# Automated and parallelized microfluidic generation of large and precisely-defined lipid nanoparticle libraries

**DOI:** 10.1101/2025.05.26.656157

**Authors:** Andrew R. Hanna, Sarah J. Shepherd, Gregory A. Datto, Isabel B. Navarro, Adele S. Ricciardi, Marshall S. Padilla, Neha Srikumar, Shuran Zhang, Hannah M. Yamagata, Nova Y. Meng, Joshua R. Buser, Michael J. Mitchell, David A. Issadore

## Abstract

Building on the success of lipid nanoparticles (LNPs) in vaccines, LNPs are being developed for a broad set of therapeutic applications by changing both the structures of the lipids used to formulate each LNP and their relative proportions. Because lipid synthesis and *in vivo* screening have been parallelized using combinatorial chemistry and LNP barcoding respectively, the manual and sequential microfluidic formulation of LNPs has become the rate-limiting step in the discovery process. In this work, we present a high-throughput, automated microfluidic platform capable of generating large, precisely-defined LNP libraries in parallel at a rate of one distinct formulation every three seconds. Each formulation is defined by varying the reagent flow ratios into one of eight microscale mixers using lithographically encoded fluidic resistors and dynamically controlled external pressure supplies. The microfluidic chip is integrated with custom frobotic plate handling for the rapid collection of each distinct formulation. Using this platform, we produce a library of 96 formulations, which we profile physicochemically and evaluate in terms of both *in vitro* and *in vivo* transfection.

## Introduction

Lipid nanoparticles (LNPs) have demonstrated broad potential for the delivery of nucleic acid cargos for gene therapy [1, 2, 3], immunotherapy [4, 5], and vaccine applications. The lipid components of an LNP formulation, namely the ionizable lipid (IL), cholesterol, phospholipid, and poly(ethylene glycol) (PEG) lipid, must be altered for each clinical application, considering desired tissue or cell tropism, immunogenicity, and other therapeutic-specific factors [6, 7, 8, 9, 10]. Though each new clinical application requires a distinct LNP formulation, there do not yet exist sufficient design principles that relate the components of an LNP and its generation method to its structure and performance *in vivo*, necessitating the screening of large libraries of formulations in biological models.

The high dimensionality of the space of possible LNP formulations has motivated researchers to develop new approaches to generate and screen LNP formulations at high throughput. Combinatorial chemistry has been used to rapidly generate diverse lipids in parallel; as a result, there now exist over 10^5^ ionizable lipids alone [11, 12, 13]. LNP barcoding can provide biodistribution data from the pooled screening of tens to hundreds of formulations in a single animal [14]. However, no LNP formulation method thus far can match the throughput of these techniques. Robotic liquid handlers can formulate LNPs at the theoretical throughput of tens to hundreds of LNP formulations per hour [15, 16] but rely on turbulent pipette mixing [17] which results in batch-to-batch variability and particle heterogeneity. Moreover, liquid handlers are typically limited to a maximum volume of 1mL per sample, requiring that a different LNP generation method must be used to formulate LNPs beyond the scale of discovery, confounding the translation of LNPs discovered using this method to animal testing and to clinical evaluation [18]. Microfluidic mixing solves the issue of heterogeneous particle formulation by rapidly mixing particle components at the micrometer scale [19, 20, 21], resulting in particles with consistent physicochemical and biological properties [8, 22, 23, 24, 20, 25]. Using recently developed microfabricated parallelized architectures such as the silicon scalable lipid nanoparticle (generation) SCALAR chip [20], microfluidic mixing can also scale over three orders of magnitude of production scale, from the discovery phase (∼ 10mL/hr) to the production phase (∼ 10L/hr) without affecting LNP properties [19, 20]. While formulating LNPs using microfluidics has been automated to some degree [26, 27], the process requires significant manual intervention and is generally serial, producing only ∼ 10 distinct formulations per hour.

To accelerate the rate of microfluidic LNP discovery, we have developed a platform for formulating distinct LNPs at a rate of 300µL of one formulation every three seconds, at least 100x faster than conventional microfluidic approaches (Figure 1). We name the platform **LI**pid nanoparticle **B**atch production via **R**obotically **I**ntegrated **S**creening (**LIBRIS**). We have designed LIBRIS using a parallelized microfluidic device architecture wherein multiple fluidic inputs each bearing an LNP component are delivered to individual LNP generators in defined proportions determined by differentially encoded fluidic resistors and dynamically controlled pressure-driven flow. LIBRIS integrates a custom-designed and microfabricated silicon-glass chip with a microcontroller-operated plate robot and pressure manifold, enabling automated control and direct collection of distinct outputs into dialysis cassettes for post-nucleation processing. To evaluate the utility of this platform, we generate a library of 96 distinct LNP formulations, comprising three sets of 32 formulations each made from a different set of lipids. We generate three sets of 32 LNPs in 90s each, characterize their physicochemical properties and transfection efficiencies, and evaluate the *in vivo* performance of a subset of the library.

**Figure 1.**
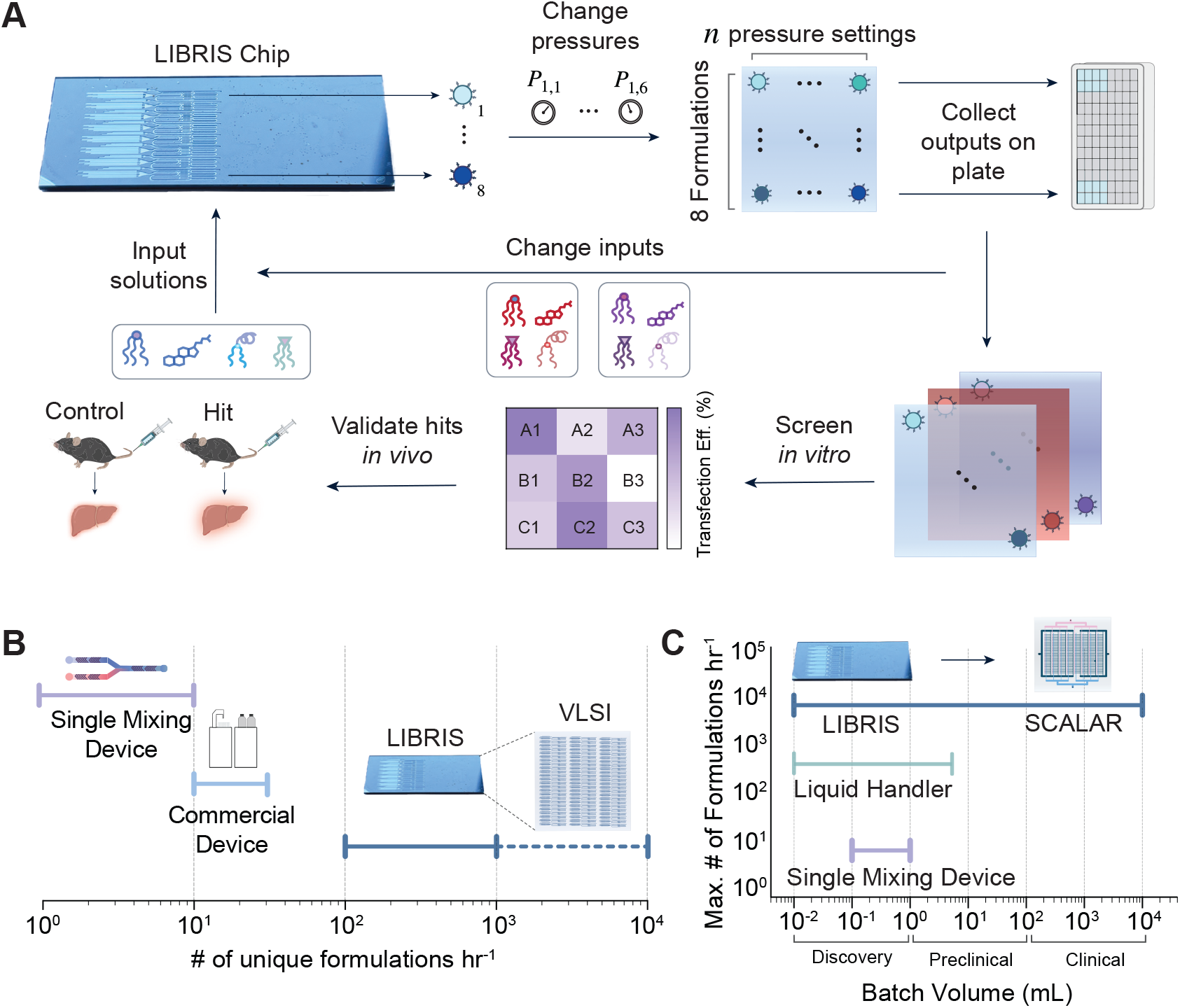
(A) Overview of the iterative generation and screening of LNP libraries using LIBRIS. (B) Comparison of the number of unique formulations per hour possible using LIBRIS and other commercially available devices for microfluidic generation of LNPs. LIBRIS could be scaled beyond 10^3^ formulations hr^−1^ by increasing the number of mixing units using very large scale integration (VLSI). (C) Depiction of the scale invariance of parallelized microfluidic mixing, demonstrated with a parallelized chip comprising many of the same mixing units and resistors (SCALAR) [20], as compared with other mixing techniques for LNP formulation.

## Results

### Integrated microfluidic LNP library generator leverages differentially encoded flow resistors across identical mixers

The LIBRIS chip comprises eight microfluidic LNP generators which mix three fluidic inputs (*i*_1_, *i*_2_, *i*_3_) containing lipid components solubilized in ethanol and three fluidic inputs (*i*_4_, *i*_5_, *i*_6_) containing RNA and citrate buffer (Figure 2A, S1). On the back side of the chip, a delivery channel distributes each fluidic input to each of the eight generators on the front side of the chip using through-silicon vias (Figure 2A, B)[28]. Upstream of each LNP generator, there are six unique lithographically defined resistors that control the relative flow rates of each of the six ethanol and aqueous inputs (Figure 2A). First, the aqueous and ethanol inlets are separately mixed in two different staggered herringbone mixers (SHMs). After separate and complete mixing of both the aqueous and ethanol inputs, they combine to formulate LNPs in the final SHM (Figure 2B).

**Figure 2.**
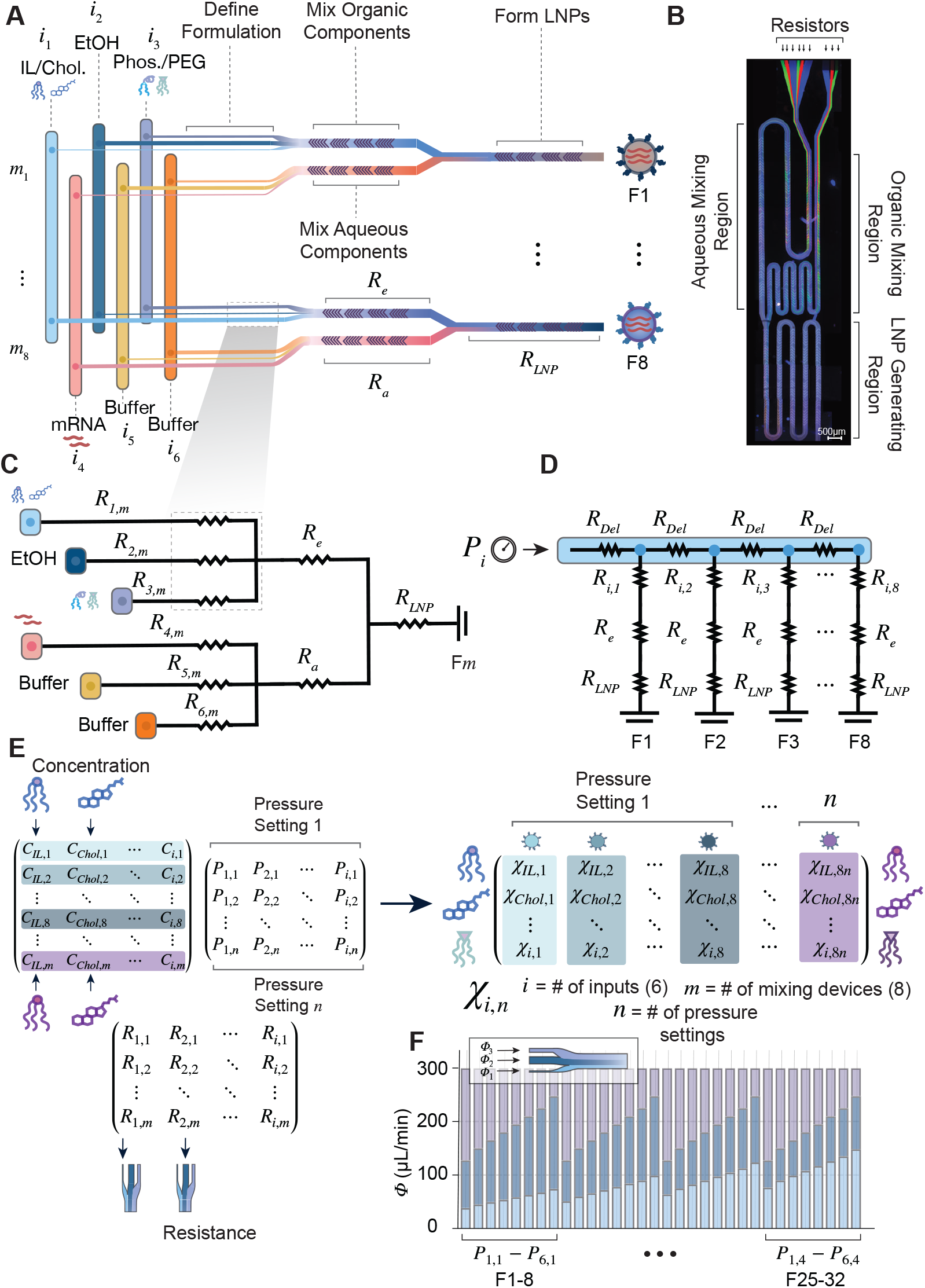
Design and fabrication of the LIBRIS chip. (A) Schematic of LIBRIS chip, depicting fluid entering one of six inputs *i*_1–6_ and being distributed differentially through resistors *R*_*i,m*_, where *i* indexes the fluidic input and *m* indexes the LNP generator. The six inputs combine in different combinations in each LNP generator *m*_1–8_ to generate distinct formulations *F* 1–*F* 8. (B) Labeled image of a single mixing unit. Circuit diagrams for the resistor network for a single (C) device and (D) fluidic input. The LIBRIS chip varies pressures *P*_*i,n*_ (E, top middle) to deliver a set of lipid structures at a given set of concentrations (E, top left) through a set of lithographically defined resistors (E, bottom), producing a library of LNPs of the size 8*n* where *n* is the number of pressure settings (E, right). (F) Though we vary relative flow rates, the total flow rate through the ethanol and aqueous mixers of each LNP generator remains the same across all pressures tested.

We designed the LIBRIS chip using a lumped-element circuit model (Figure 2C–D, S2, S3). Based on previous approaches [19, 20, 28], we incorporate lithographically defined microfluidic flow resistors upstream of each LNP generator (Figure 2C), where the resistance of each resistor *R*_*i,m*_ is much greater than the resistance of the downstream SHMs such that the total resistance of each LNP generating device is approximately *R*_*i,m*_, for each fluidic inlet *i* and LNP generator *m*. We model the fluidic resistance of each element in LIBRIS, assuming a rectangular cross section of dimensions length *L*, width *w*, and height *h* (assuming here that *h* is the smallest dimension) using the relation [29]

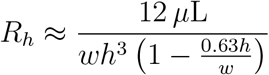

We vary the resistance of each of the fluidic resistors *R*_*i,m*_ by varying the width of each resistor. Modifying the height across different devices would require many additional lithography steps and would be more sensitive to fabrication errors due to the strong non-linear dependence of *R*_*i,m*_ to the limiting dimension *h*. We maintain the same length for each resistor to simplify connection of each generator to the delivery channels.

We model our chip architecture as a ladder geometry where the fluidic resistance of the delivery channel between each of the generators *R*_*Del*_≪*R*_*i,m*_ (Figure 2D, Figure S2) [30]. Thus, we assume that the flow rate *ϕ*_*i,m*_ through each resistor is defined by the pressure applied to each inlet *P*_*i*_ divided by the resistance of each resistor *R*_*i,m*_. We validate this assumption by simulating the circuit across the range of our operating pressures and performing a pairwise distance analysis between the flow rates through each generator generated by the circuit simulation and by the values calculated using the assumptions above (Figure S3). We observe a strong correlation of simulated and calculated relative flow rates across all LNP generators (*R*^2^ = 0.99) (Figure S3C) as well as simulated and actual relative flow rates (*R*^2^ = 0.97) (Figure S3D–E). Thus, with an integrated pressure manifold, we can rapidly modulate each inlet pressure *P*_*i*_ to vary the relative flow rate of the LNP components through the resistors *R*_*i,m*_ to generate many distinct sets of eight formulations (Figure 2E).

We designed the LIBRIS chip to maintain a constant total flow rate through each generator, even as the relative flow rates of LNP components vary, allowing the lipid stoichiometry to change while preserving consistent mixing conditions (Figure 2F, Figure S2). We achieve this function by using one of the aqueous and ethanol inputs to compensate for the flow rates from the other inputs are varied. We designed the flow rates of each ethanol input (*i*_1_, *i*_2_, *i*_3_) at a 1:3 ratio to their corresponding aqueous input (*i*_4_, *i*_5_, *i*_6_) to maintain the same ratio of IL from *i*_1_ and mRNA from *i*_4_ across all LNP generators [13, 31, 32] and to maintain the total flow rate ratio of ethanol and aqueous inputs at 1:3, as is commonly done in the literature [23].

### Silicon and glass chip architecture permits highly precise featuring and chemical inertness at high pressures

We fabricated the chip by photolithographically patterning and then anisotropically etching four unique layers into a single 500µm thick, 100mm diameter silicon wafer that we anodically bond to glass wafers, adapting a fabrication strategy previously published by our group (Figure S4–6) [20, 28]. We chose to use silicon and glass for their capacity to be patterned with sub-micron resolution [33], to be operated above 1000 psi [34], and to be compatible with a wide variety of solvents [35]. Choosing to manufacture a device with features as small as those of our fluidic resistors in soft materials, such as PDMS, would result in uncertainty in flow rates due to channel deformation at our typical operating pressures [36]. After fabrication, chips were diced from finished wafers and then housed inside a custom aluminum manifold, machined by Protolabs™ with inlet ports and outlet ports spaced to fit a 96-well plate (Figure S7–8).

### Characterization of the LIBRIS chip reveals efficient and uniform mixing across all LNP generators and validates intended flow rate ratios

We characterized the mixing efficiency of each of the eight LNP generators and determined both that their mixing rates were the same and that the relative flow rates through each mixer were as designed (Figure 3). To perform this characterization, we used solutions of three water-soluble fluorescent dyes with minimal spectral overlap as each of the ethanol inputs (*i*_1_, *i*_2_, *i*_3_) and the corresponding aqueous inputs (*i*_4_, *i*_5_, *i*_6_). While operating the chip at the pressure settings later used to generate LNP libraries (Table S1), we imaged the chip using a fluorescence microscope, ensuring that intensity was within the linear dynamic range of the camera across all fluorescence channels (Figure 3B) [20, 25].

**Figure 3.**
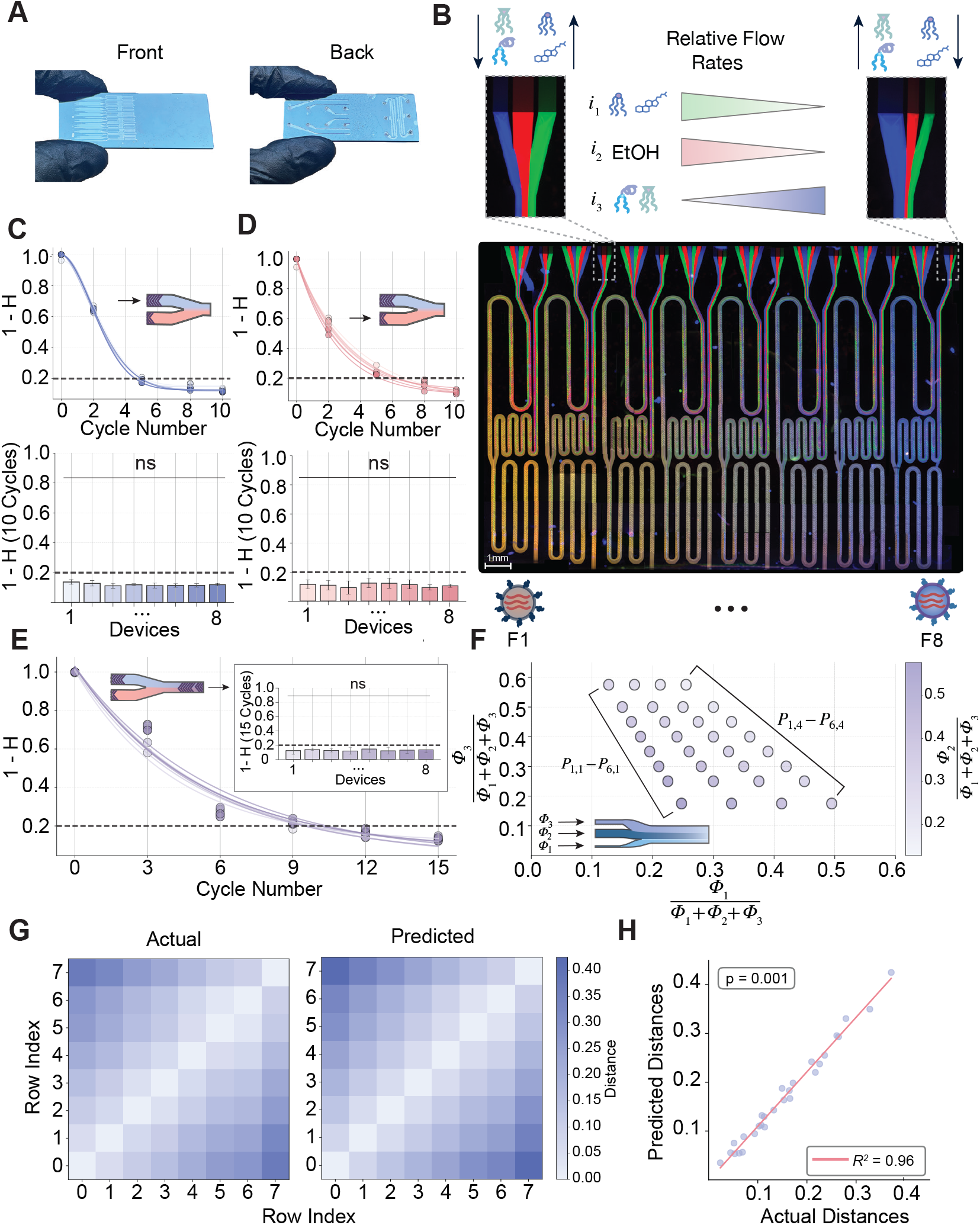
Characterization of mixing efficiency and relative flow rates across LIBRIS chip generators. (A) Image of the fabricated LIBRIS chip. (B) Fluorescent dyes were imaged flowing through the device. *i*_1_ is schematized as containing the IL and cholesterol, while *i*_3_ contains phospholipid and PEG lipid. Mixing efficiencies were quantitated (±SD; errors determined by fitting 1 − *H* to an exponential decay model) and compared using a one-way ANOVA followed by Tukey’s HSD post-hoc test across the (C) ethanol (*F* = 0.001, *P* = 1.0), (D) aqueous (*F* = 0.006, *P* = 1.0), and (E) combined mixing regions of the chip (*F* = 0.004, *P* = 1.0). (F) The assumed flow rate ratios for each mixing device were compared using (G) pairwise distance analysis to the actual flow rate ratios. (H) Correlation between pairwise distance matrices was compared using a Mantel test (*R*^2^ = 0.96, *P* = 0.001).

To quantify the efficiency of mixing (*H*) for each device, a fluorescence micrograph was taken of each mixing portion, and three fluorescence channels were overlaid. To eliminate non-specific background, greyscale values of a brightfield image of the corresponding mixing portion were subtracted, and then the mean squared deviation of the dye intensity for each dye was integrated with respect to position *y* across the width *w* of the channel [37, 38]. The mixing efficiency was calculated from the fluorescence intensity values *I* at each pixel as follows [38]:

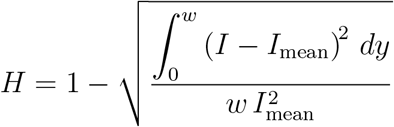

We calculate *H* for each of the three dyes used and report the arithmetic mean of all three values. When 1 −*H* is plotted against the number of mixing cycles, the data fits an exponential decay model (Figure 2C–E). Consistent with previous literature, we observe a plateau in mixing efficiency before the end of each of our SHM regions — 10 mixing cycles for the aqueous and ethanol SHMs and 15 mixing cycles for the LNP generating SHM. Consistent with previous literature, we consider complete sample mixing at 1−*H*≈ 0.2 [38]. Within each group of ethanol, aqueous (Figure 3C–D), and LNP generating SHMs (Figure 3E), we observe the mixing value of 1 − *H <* 0.2 at the same mixing lengths, indicating that observed differences in LNP characteristics derive from differences in lipid ratios between generators, not differences in mixing efficiency.

From the same mixing experiments, we evaluated the relative mixing ratios of the different LNP components through each LNP generator. We chose the resistances *R*_*i,m*_ through each of the three ethanol phase inputs *i* of the LIBRIS chip to follow the relation:

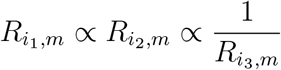

By doing so, the concentration of components can vary either directly or inversely proportionally across the eight generators depending on which inputs bear the components. We observed this relation qualitatively in the color gradient of the generator outputs across the chip, where the flow rates of green dye (*ϕ*_*i,*1_, ϕ_*i,*4_) and red dye (*ϕ*_*i,*2_, ϕ_*i,*5_) flow rates trend inversely to the blue dye (*ϕ*_*i,*3_, ϕ_*i,*6_) flow rate across the generators (Figure 3B, Figure S2).

To quantify the relative ratios of each fluidic input in each mixer, we subtracted the greyscale values of a brightfield image of each fluorescence micrograph, normalized the intensity of each dye relative to an unmixed portion of the device, and calculated the intensity ratio of each dye between the fully mixed and unmixed portions of the device. We opted to measure relative intensities rather than to measure laminar thicknesses given the effects of boundary conditions on the fluidic velocity profile in a channel of rectangular cross section [29]. To compare the difference between the predicted and actual flow rate ratios of the LIBRIS chip, we normalize flow rates through each mixing unit at each pressure setting and perform a pairwise distance analysis on the relative flow rates through each input (Figure 3F, inset) of our calculated distribution (Figure 2F, 3F) and actual distribution (Figure 3G) as determined above, observing that the distributions of relative flow rates are nearly identical as compared by a Mantel test (*R*^2^ = 0.96, *P* = 0.001) (Figure 3H).

### Design and implementation of custom plate robotic interface allows for rapid collection of distinct LNPs to in-line dialysis

Though the LIBRIS chip rapidly generates many formulations from a single set of solutions simply by varying pressures, each of these outputs must be kept separate for further screening. Thus, we designed and fabricated an interface that coordinates the motion of a custom plate robot with the control of a set of programmable pneumatic regulators, enabling disposal of undesired priming solution and collection of pure sample directly into on- plate dialysis cassettes once the chip has equilibrated (Figure 4A–B).

**Figure 4.**
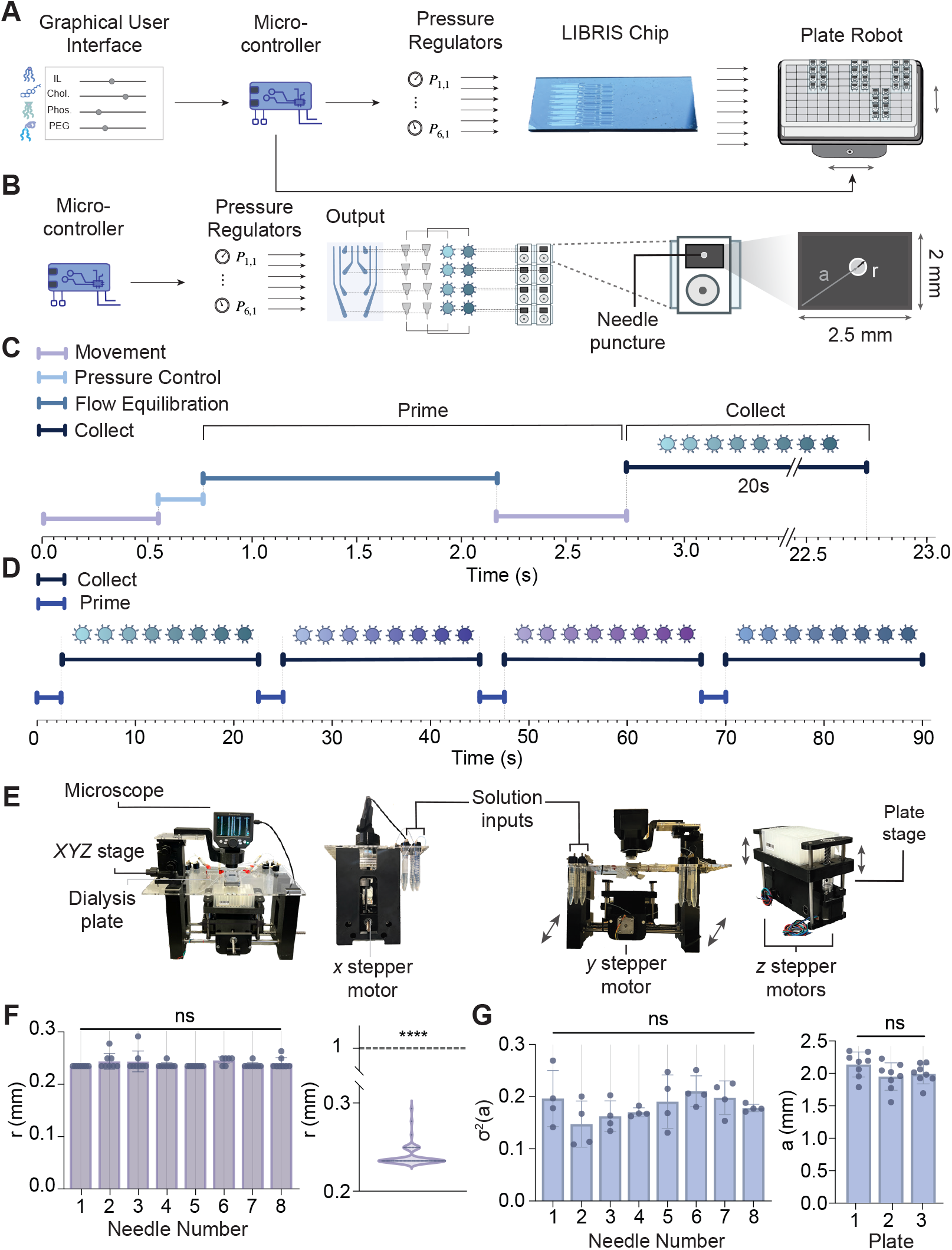
Schematic overview, timing diagram, and collection consistency of custom plate robotic interface. (A) Overview of a single collection run, where a user initiates the setup and the microcontroller synchronizes plate motion with pressure regulation. (B) The life cycle of a single set of eight LNPs generated and collected by the plate robotic interface. After initiating, (C) a timing diagram shows the breakdown of the 2.6s waste cycle for a set of eight LNPs, (D) where a total of 90s is required to collect 32 LNPs. E) A labelled set of images of the plate robot. Measurement of the precision of collection in F) *n* = 20 iterations of movement on the same plate and (G) *n* = 20 iterations of movement across three separate plates. (F, left) Measurements of puncture marks across four collection positions for the same needle position were compared using a one-way ANOVA followed by a Tukey HSD post-hoc test (*F* = 1.5, *P* = 0.19). (F, right) A one-sample t-test was performed to determine whether the radius of puncture was below the acceptable radius of 1mm (*t* = − 572.4, **** = *P <<* 0.0001). (G) The variance of the puncture marks at the same positions across three plates (left) were compared similarly (*F* = 1.4, *P* = 0.27) and absolute position for each needle within the cassette was compared across plates (right, *F* = 2.2, *P* = 0.14).

The step-by-step process for LIBRIS to generate a set of distinct formulations is as follows: it initialize its flow under a new set of pressures, discards the output associated with this initialization, moves the plate to collect the eight outputs into individual dialysis cassettes, and collects ∼ 300µL of each formulation, which takes around 22.5s, i.e. 2.75s per formulation (Figure 4C–D). In this time per formulation, the integrated robotic system both equilibrates the chip and moves between each 4 × 2 array of dialysis cassettes to collect sets of eight samples. By integrating direct serial control of the regulators, the microcontroller initiates the pressure regulators in under 150ms, and subsequent equilibration of the pressure regulators occurs within 1.5s, as dictated by the proportional-integral-derivative (PID) controller (Figure S9, Video S1). From the time of pressure initiation, stabilization of the fluidic lamina on chip occurs in 1.6s, determined by analyzing the laminar flow versus time using inlets that contain fluorescent dye. To minimize the time it takes for the robot to move the chip to the next set of wells, motion of the plate from one position to another begins 600ms before the chip has finished equilibrating flow and enters the collection well at 2.75s (Figure 4C, Video S1). We achieved these exchange times between samples by minimizing on chip volume, minimizing tubing length and volumes, incorporating a silicon and glass architecture to improve the response time of the fluid flow to changes in pressure relative to mechanically softer devices made of soft polymers, and designing the plate robot to be as stiff as possible to improve its mechanical response time. The highest total dead volume of any output is ∼ 35µL, comprising the volume of the chip outlets and outfitted needle (Figure S8). Thus, at a flow rate of 20µL/s, the dead volume of each output has been completely flushed between collections. The total collection time for 32 LNPs is ∼ 90s, with *>*10s of waste collection (Figure 4D).

We measured the precision of the positioning of the plate beneath the fixed LIBRIS chip to ensure that no leaking or cross-contamination of the outputs would occur. Each chip output is connected to a needle which we position above the 2mm × 2.5mm aperture of a dialysis cassette situated in a 96-well plate (Figure 4B, S8, S10–11). By covering each cassette with a disposable membrane, we prevent any backflow or leakage from the cassette and collect the sample directly into dialysis. We measured the precision of the plate positioning by programming the robot to move through four collection positions 20 times in a row on the same 96-well plate and measuring the radius *r* of the puncture marks made by the collection needles (Figure 4F, S10–S11). The diameter of the puncture marks across all cassettes measured only 130µm greater than the diameter of the needle, meaning that any centered needle is exceedingly unlikely to puncture outside of the acceptable radius of 1mm from the center of the cassette (Figure 4F). We further evaluated plate positioning across multiple plates by measuring the puncture marks relative to a fixed position (Figure 4G, S10–S11). No significant difference was observed across the variance of puncture marks at the same movement position across three plates (Figure 4G).

We then evaluated the consistency of total sample volume collection between wells and the minimum collection volume possible to determine the versatility of the LIBRIS platform across volumes. Using 10s collection times, consistent sample volumes of 120*µ*L were collected across all outputs (Figure S12A), and the platform was capable of collecting a minimum volume of 10*µ*L per sample, enabling screening of physicochemical characteristics with minimal lipid and RNA use (Figure S12B).

To improve the accessibility of our robotic system, we designed a userfriendly graphical user interface (GUI) which converts a set of input concentrations and a desired range of molar percentages for the LNP components into the correct input pressures (Figure S13). Since we designed and printed all parts except the motors and various structural components such as support rods and bearings (Figure 4F), the total cost for the single prototype robotic system is under $500 and would likely cost significantly less if produced at scale.

### Physicochemical characterization of 96 LNP library validates production of distinct LNPs

To validate the creation of LNPs with different physicochemical properties using the LIBRIS platform, we formulated a library of 96 LNPs. We refer to each group by shorthand for the IL – D-Lin-MC3-DMA (MC3), SM-102, and C12-200. We chose to use the MC3 and SM-102 given their clinical validation and to use C12-200 given its status as a gold standard for transfection efficiency [6, 39, 40]. The design space assayed during these experiments varies the molar percentages of IL and cholesterol in direct proportion, which both vary inversely with the concentration of phospholipid and PEG lipid (Figure 5A). Using the LIBRIS chip, we formulate each of these lipids into a set of 32 LNPs at the molar ratios outlined (Table S2-4), as determined by the stock concentrations and the relative flow rate ratios derived from the mixing experiments above. We measured the hydrodynamic diameter (size) and polydispersity index (PDI) of each sample using dynamic light scattering, observing distinct sizes which correlate inversely with the percentage of PEG both within each set of eight LNPs and across all groups (Figure 5B, insets) [16, 41]. Across all samples, only three formulations demonstrated a polydispersity index (PDI) above 0.3 and only one formulation demonstrated an encapsulation efficiency (EE%) of lower than 70%, meaning that nearly all particles fall within acceptable physicochemical benchmarks (Figure 5C, D). Unlike size, EE% and PDI did not correlate with the molar concentration of PEG (Figure S14).

**Figure 5.**
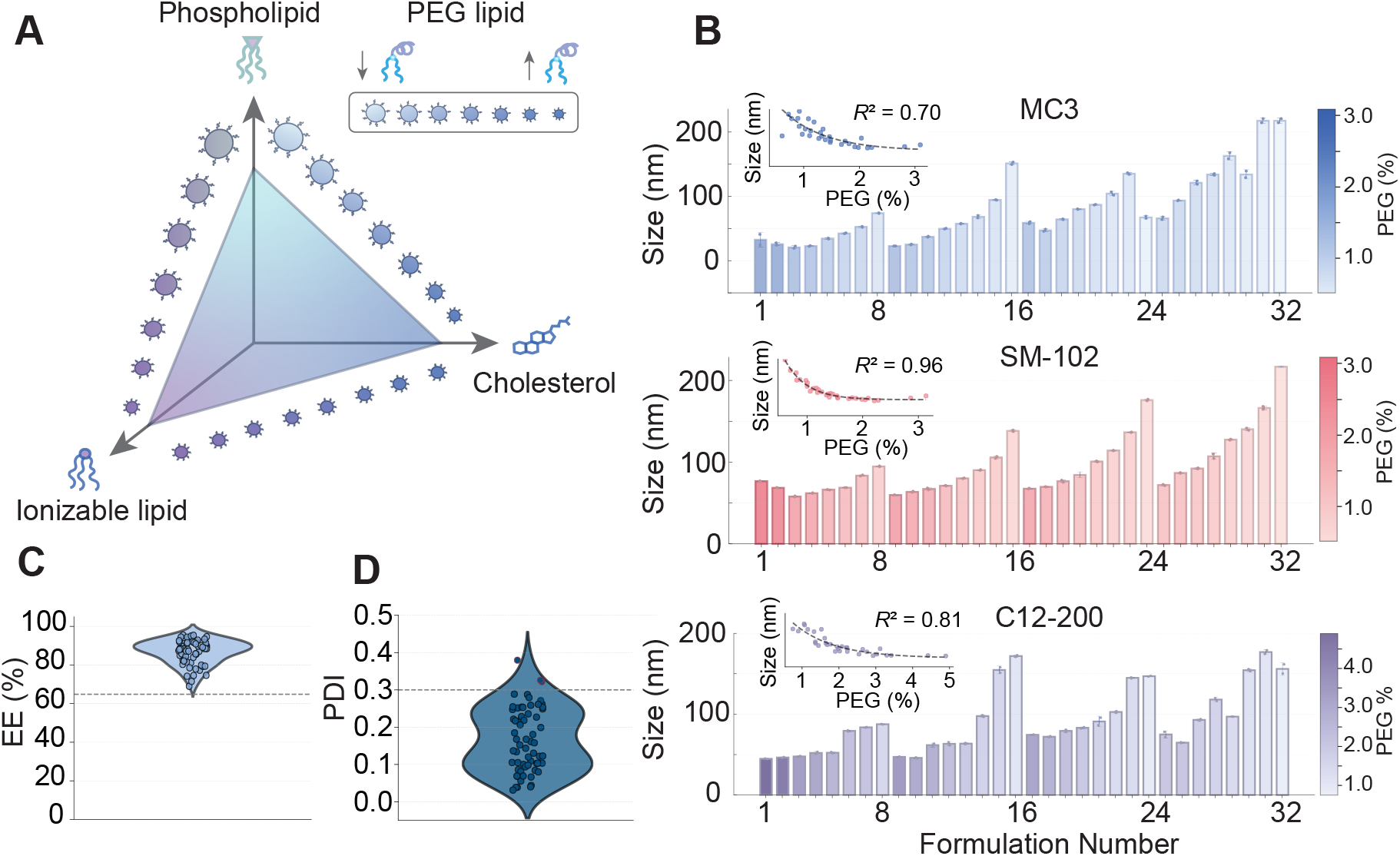
Physicochemical characterization of the LNP libraries generated by the LIBRIS chip. (A) Schematic of the potential design space assayed in this library. Across each lipid set, IL and cholesterol mole percent ratios vary directly proportionally with one another, which vary inversely with the phospholipid and PEG lipid ratios. A decrease in LNP size is shown with increased PEG concentration, schematized in the top right of the subpanel. (B) DLS measurements of the size of each formulation. (B, insets) Size decays exponentially with the molar percentage of PEG (MC3: *P <* 0.0001, SM-102: *P <* 0.0001, C12-200: *P <* 0.0001). (C, D) 95*/*96 particles and 93*/*96 particles fall within physicochemical benchmarks of encapsulation efficiency (EE%) and polydispersity index (PDI).

To ensure that the differences in physicochemical characteristics between sets of eight LNPs were not simply an artifact of drift between samples, we continuously operated the chip to generate the same formulation 5 times. Across a representative sample of three SM-102 formulations, we measured consistent particle size between collections at the beginning and end of operation, indicating that particle characteristics do not meaningfully change within the operating time of the experiments performed (Figure S15). We further compared the physicochemical characteristics of LNPs generated using our SHM-based mixing technique against automated pipettes, meant to mimic conventional liquid handlers. We demonstrate that using the LIBRIS chip, we generate significantly smaller particles (∼ 80nm vs. ∼ 180nm) with higher batch-to-batch consistency and lower, more consistent PDI than when using automated liquid handling (Figure S16). Additionally, we showed that LNPs made using the LIBRIS platform compare in size and PDI to particles made using a control PDMS SHM, and a 10× parallelized SHM (Figure S15). Thus, as demonstrated here and by our previous work [20], we can screen for an optimal LNP formulation at discovery scale, then generate a formulation with identical properties using the same mixing architecture at market scale.

### *In vitro* and *in vivo* performance of LNPs from library validates differences observed in physicochemical characterization

We evaluated the utility of the LIBRIS chip in the LNP screening process by measuring the transfection efficiency of particles across the previously described libraries (Figure 6A). Using RNA concentrations determined during EE% characterization of the libraries, we dosed HepG2 cells with LNPs loaded with mCherry from the MC3, SM-102, and C12-200 groups (Figure 6A). Using flow cytometry, we determined both the percentage of mCherry+ cells and the mean fluorescence intensity (MFI) for n = 5 wells, where we observed difference in LNP transfection efficiency contingent on formulation (Figure 6A, S17–18). To determine which of the physicochemical characteristics had the most meaningful implications on transfection efficiency, we performed a principal component analysis, excluding the LNPs which failed to meet previous physicochemical benchmarks (Figure S19A). We observed that, excluding compositional variables, which varied in fixed increments between samples, either EE%, seen in the MC3 group, or size, seen in the SM-102 and C12-200 groups, had the most significant loading across the groups, a result generally consistent with the literature [42, 43]. To analyze the relationship between size – a variable with significant loading across all samples – and transfection efficiency, we performed Pearson correlation analyses between transfection efficiency and PEG lipid molar composition across each group, revealing that in the MC3 and SM-102 groups, PEG percentage and transfection efficiency correlate inversely, while in the C12-200 group, transfection efficiency and PEG percentage correlate directly (Figure S19B). To evaluate the relation between the *in vitro* performance of particles made using our chip and their *in vivo* performance, we quantified the *in vivo* hepatic transfection from the top performing LNP (C12-200 F4), the poorest performer (C12-200 F24), and the previously optimized C12-200 formulation (Figure 6B, S20) [6]. Instead of formulating these using the LIBRIS chip, we sought to cross-validate the results of the *in vitro* screen by determining the molar concentrations of each lipid corresponding to the formulations above (Table 1) then formulating each LNP using a single microfluidic mixing device as described previously [19, 44]. Separate cohorts of mice were treated intravenously (i.v.) with each of these formulations loaded with firefly luciferase mRNA at a total dose of 0.1mg/kg (Figure 6C). Expression was observed almost entirely in the liver, consistent with previous literature (Figure 6C, Figure S21) [6]. Total hepatic luminescence indicated around a two-fold increase in expression between F4 and F24, with F4 performing comparably to the previously optimized LNP formulation (Figure 6C) [45, 46]. Hepatic luciferase expression is reported here since it significantly outweighed the expression of all other organs similarly across all groups (Figure S21). These results further validate the differences in composition generated in each library by the LIBRIS chip and suggest that there exist more biologically optimal LNPs discoverable by varying compositional ratios within a single set of lipid structures.

**Table 1:** Physicochemical characteristics and molar compositions of LNPs tested *in vivo*.

| Formulation | IL (%) | Cholesterol (%) | Phospholipid (%) | PEG lipid (%) | Size (nm) | PDI |
| --- | --- | --- | --- | --- | --- | --- |
| F4 | 33.3% | 44.3% | 19.4% | 3.0% | 73.8 | 0.20 |
| F24 | 40.1% | 53.3% | 5.7% | 0.9% | 121.8 | 0.06 |
| Base Formulation | 35% | 46.5% | 16% | 2.5% | 63.8 | 0.26 |

**Figure 6.**
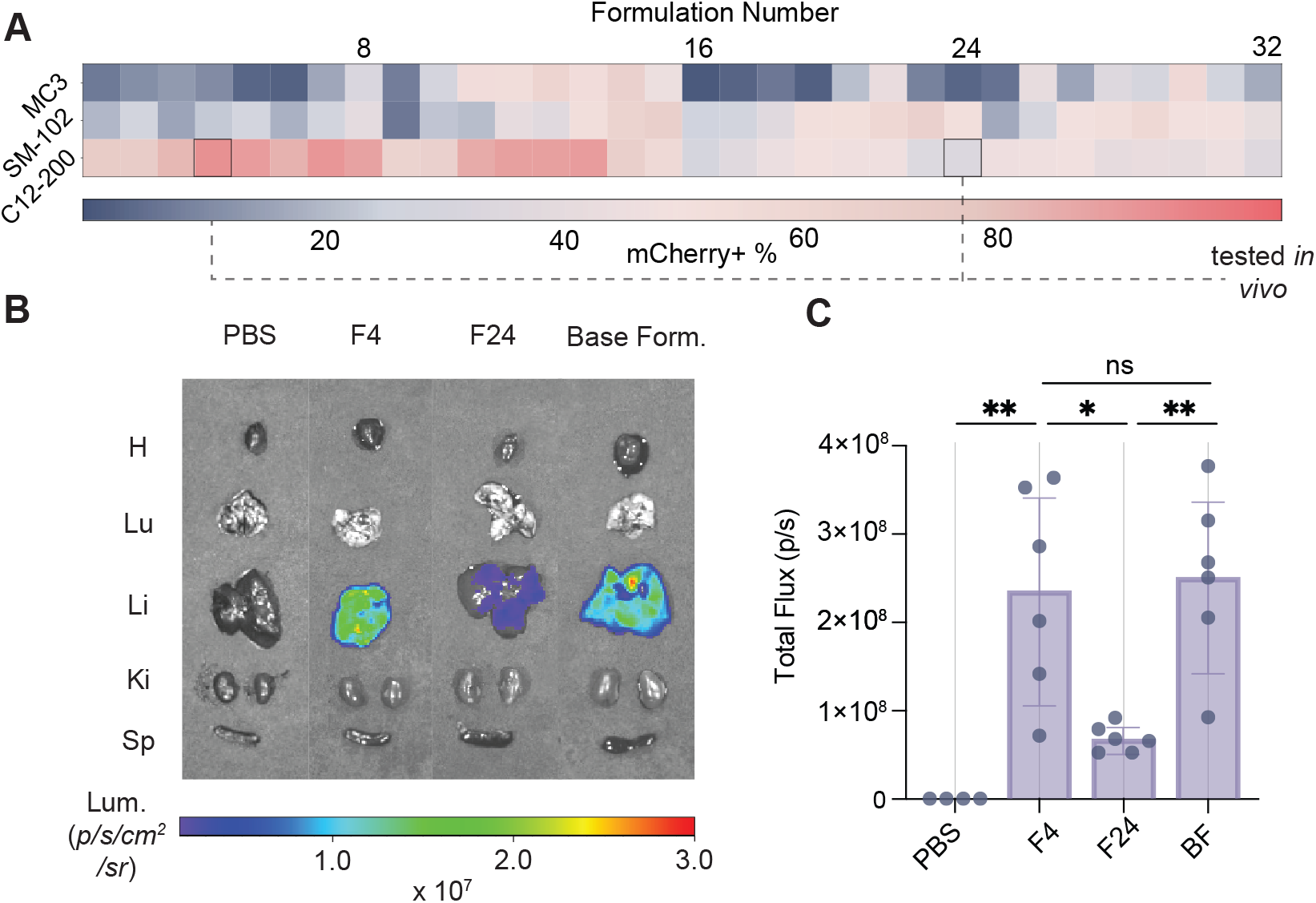
*In vitro* and *in vivo* validation of differences between LNPs in libraries shown by physicochemical characterization. (A) Heat map of the % of mCherry^+^ HepG2 cells across all 96 LNPs screened. The lowest and highest transfecting C12-200 LNP moved forward from the *in vitro* screen into *in vivo* testing. (B) Representative IVIS images of the major organs extracted from one member of each cohort 6hr after LNP administration showing luciferase expression in the liver (H, heart; Lu, lungs; Li, liver; Sp, spleen; Ki, kidneys). (C) Quantification and analysis of hepatic luciferase expression between each treatment group. Normality of each sample distribution was validated by a Shapiro-Wilk test (PBS: *P* = 0.5881, *F* 4: *P* = 0.5913, *F* 24: *P* = 0.3048, BF: *P* = 0.6586). One-way ANOVA followed by Tukey’s HSD test was used to compare total luminescence between all groups (\**P <* 0.05, \*\**P <* 0.01). *n* = 4 mice in PBS group. *n* = 6 in treatment groups.

## Discussion

We present an approach that significantly enhances the throughput of generating precise microfluidic formulations of distinct LNP formulations. By parallelizing the delivery of lipid components on our LIBRIS platform through lithographically encoded resistors, we achieve at least a 100-fold increase in throughput relative to manual microfluidic device operation. The LIBRIS platform has the potential to accelerate the discovery pipeline over the state of the art by enabling much more rapid and precise screening of formulation spaces for existing or previously untested sets of lipids.

To further improve the throughput of LIBRIS, it is possible to increase the number of devices 10× or 100× on the space of a silicon wafer up to 6” or 9” in diameter. We have previously expanded the design principles of a ladder geometry of parallelized resistors fed using delivery channels outlined here to 20,000 generators on a single chip for droplet generation [28]. While the current dynamic range between the lowest and highest flow rate of each input to a LNP generator is around 1 : 4.5, we could improve this dynamic range by increasing the resistance of each input, preventing back-flow and providing the ability to generate larger compositional differences at any one set of pressures. Ultimately, even if we could generate many more distinct formulations per pressure setting on chip, the collection of distinct formulations in millimeter scale compartments for later screening and handling remains a limiting challenge [47, 48]. Future innovations for on-chip separation and labeling of distinct formulations or off-chip particle collection will be necessary to fully leverage the benefits of on-chip parallelization.

Beyond the improvement in throughput over manual and sequential formulation of LNPs, the LIBRIS platform can produce formulations with less waste and with greater control over sample volume. Given the parallelized and automated nature of the platform, the waste volume per formulation is on the order of 10× less than generating these formulations manually, meaning that smaller amounts of costly RNA and lipids can be used to generate each formulation. The LIBRIS platform can be used to generate volumes as low as 10µL of many distinct formulations for screening, then using the SCALAR chip [20], manufacturing of that formulation can be scaled to L/hr under the same mixing conditions. Additionally, since the chip is manufactured out of silicon and glass, it can be rapidly cleaned at high pressures [20] or modified with antifouling coatings [49, 50]. Because our robot measures around 18” × 18” ×10” (*l* ×*h* ×*w*), the entire setup can be situated in a biosafety cabinet to generate particles under sterile conditions. The ability of LIBRIS to generate large, precisely-defined LNP libraries is an advancement toward generating the data sets necessary to train machine learning models that can better predict the physicochemical and biological properties of new drug candidates for specific applications. While there exist some successful predictive models for new LNP formulations to train data-intensive deep learning models [51, 52, 15, 53], these models require larger data sets across a greater number of biological models. Future work will focus on coupling LIBRIS to additional automation to interchange lipids, allowing for the fully automated microfluidic formulation of many sets of combinatorially generated lipids across more biological models.

The LIBRIS platform addresses the current rate-limiting step in the consistent generation of large LNP libraries for screening and provides a path forward for a more rigorous evaluation of a larger subset of the potential compositional ratios for a given set of lipid structures, enabling the future use of large scale machine learning models for the development of function-optimized LNPs.

## Methods

### 0.1 Fabrication of LIBRIS chip

Chips were fabricated in the Quattrone Nanofabrication Facility at the University of Pennsylvania. The chip design comprises four layers, designed in AutoCAD (San Rafael, CA): the mixing channels (Layer 1, depth: 85µm), the resistors and herringbones (Layer 2, depth: 20µm), the delivery channels (Layer 3, depth: 365µm), and the “through silicon vias” or interconnects between both sides of the wafer (Layer 4). In brief, using a Heidelberg 66+ laser writer (Heidelberg, Germany), chrome-coated soda lime photomasks (Telic Company, Santa Clarita, CA) were patterned, developed in AZ300 MIF developer (EMD Performance Materials Corp., Philadelphia, PA), and then the exposed pattern was etched into the chrome (Transene Company, Danvers, MA). Remaining photoresist was removed by PE Remover (DuPont, Wilmington, DE) under sonication at held at 65 °C for 10 min. The general pattern for fabrication of each layer etched into a single 500µm thick, 100mm diameter silicon wafer (ID 775; University Wafer, South Boston, MA) is as follows: prepare wafer with resist adhesion promoter, spray coat resist, bake, rest, lithographically pattern, rest, develop, etch, perform metrology, adjust etching if feature depths are incorrect, and clean before moving to the next layer. In greater detail, at the beginning of each layer, the wafer was coated with 49% HF for 30s, then rinsed thoroughly with water, and dried on a 200°C hotplate for 2min. S1805 photoresist (Dow, Midland, MI) was mixed at a 1:8 ratio with acetone, then spray-coated on the processed wafer to desired thickness using an AS8 AltaSpray Coater (SÜ SS MicroTec, Garching, Germany). Wafers were baked at 110°C for approximately 5min per 4µm of resist immediately after spray coating, and then left to rest at room temperature for ¿30min for each step to prevent outgassing during exposure. The respective resist thicknesses for each layer 1–4 are 8µm, 8µm, 16µm, and 4µm. After spray coating, resting, and baking, a mask aligner (MA6, SÜ SS Microtec) was used to expose the coated wafer. Exposure times varied depending on resist thickness. After exposure, each wafer was developed in AZ300 MIF developer, then rinsed, dried at room temperature, and subsequently underwent a deep reactive ion etch (DRIE) (SPTS Rapier Si, Newport, UK) to the etch depths listed above. After etching, each wafer was profiled using an optical profilometer (Profilm 3D, Filmetrics, San Diego, CA), and cleaned using PE remover and Nano-Strip (VWR, Radnor, PA) heated to 110°C. This process was repeated for each layer.

Layer 2 (resistors and herringbones) underwent a slight modification to the procedure to preserve the edges of the mixing channels where resist can delaminate during the exposure process. After cleaning from the end of Layer 1, the wafers were conformally coated with 500nm of SiO_2_ using plasma-enhanced chemical vapor deposition (Oxford Plasma Lab 100 PECVD) (Oxford Instruments, Abingdon, UK). The layer process proceeded as described until after exposure, where the exposed 500nm layer of SiO_2_ was etched away using CF_4_ plasma in the reactive ion etcher (Oxford 80 Plus, Oxford Instruments, Concord, MA).

For Layer 4, a 4µm thick layer of SiO_2_ was deposited using PECVD on the the delivery channel side (backside) of the wafer to prevent the breaking of vacuum lock under the final through silicon via DRIE step. Instead of promoting resist adhesion using 49% HF, which would rapidly dissociate this oxide layer, the wafer was coated with bis(trimethylsilyl)amine (HMDS) (Genesis Prime Oven, Genesis Systems, Davenport, IA) for 10min.

The fully etched wafer, and two 100mm Borofloat 33 glass wafers (ID 517; University Wafer), one of which was micromachined with a custom inlet-outlet hole setup using an IX-255 laser system (IPG Photonics, Oxford, MA), are left in Nano-Strip at 110°C for 30 min. All wafers are spin-rinse dried (RENA Compass, RENA, Albany, OR), then stacked in a glass/silicon/glass arrangement, aligning the inlet-outlet glass holes with the etched delivery channels. The triple stacked wafer is anodically bonded at 900V and 400°C under 100N piston force using an EVG 510 Wafer Bonding System (EVG Group, Oberosterreich, Austria), then the wafer stack is let to cool to at least 180°C before flipping the wafer stack over to repeat the bonding in the other direction, producing a bonded triple stack of wafers. The wafers are diced into single chips using a dicing saw (ADT 7100, ADT, Horsham, PA) fitted with a resin blade.

### 0.2 Circuit simulation

A replica of the chip was rendered in LTSpice Version 17.2.4. Pressure (Pa) was analogized as voltage for each respective fluidic inlet *i* across each set of pressures that we experimentally validated 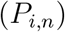.Flow rate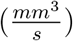 was analogized to current. Hydraulic or fluidic resistance 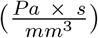 was analogized to electrical resistance. Hydraulic resistances were calculated from dimensions determined by measurements of each chip using optical profilometry (Profilm 3D). Pairwise distance analyses were performed between relative flow rate values derived from the LTSpice simulation and both predicted values and actual values. Correlation coefficients were calculated using a Mantel test (Figure S2).

### 0.3 Mechanical integration of pneumatic regulator setup with chip

A custom housing for the chip was designed in Fusion360 (Autodesk) and machined from 6061-T651 aluminum (Protolabs, Maple Plain, MN). 1/4” – 28 UNF threaded ports were included on the backside of the chip to allow for collection of inlets and outlets and viewing windows exposing all mixing channels were located on the front side of the chip (Figure S8). A nitrogen tank was connected to six dual-valve pressure controllers (Alicat Scientific, Tucson, AZ), via a 7-port manifold for 1/16” outer diameter (OD) tubing (IDEX H&S, Oak Harbor, WA). Each regulator was connected to a pressurizable vessel (Elveflow, Paris, FR) via flat bottom 1/4”-28 PEEK M-UNF fittings (IDEX H&S). These vessels contained the fluidic inputs, which were distributed to the separate delivery channels on chip through a pressure fit connection between the machined glass and a 1/16” OD ferrule (IDEX H&S). For the collection of outputs, eight M-UNF 1/4”-28 x Luer slip adapters (IDEX H&S) were threaded into the 1/4”-28 M-UNF output ports of the aluminum housing, and 30G x 9mm low-dead volume needles (Air-Tite, Virginia Beach, VA) were press fitted to the adapters. An additional manifold was cut from 1/8”-thick clear acrylic and fit over the collars of the needles to better align and support the needles in the 4 × 2 array.

### 0.4 Design and fabrication of the plate robot

All parts of the skeleton of the robot, including robot legs, plate stage, and motor compartments, were designed in Fusion360. Lead screw nuts were printed using a Form2 SLA printer with Durable Resin (Formlabs, Somerville, MA). All other components were printed from PolyLite™ PETG filament (Polymaker, Shanghai, China). The top base plate which held both the chip and the *xyz* stage for the portable microscope was laser cut from 1/4” thick clear scratch-proof acrylic (McMaster-Carr, Robbinsville, NJ) with a 0.0075” tolerance. Additional parts, such as stainless steel and ceramic-coated 6061-T651 aluminum shafts (3/8” Diameter, 9” Long), lead screws (1018 Carbon Steel Precision Acme Lead Screw, Fast-Travel, Right-Hand, 3/8”-5 Thread, 5:1 Speed), and stainless steel bearings (3/8” Bore, 3/8”×5/8” ×1-19/32”) were acquired (McMaster-Carr), and the skeleton of the robot was fabricated using additional bolts, nuts, and spacers from the GM Lab at the University of Pennsylvania.

The skeleton of the robot was fabricated by first assembling the central plate support as shown (Figure 4E). This assembly was then mounted on two 12-inch linear motion shafts (Tapped Linear Motion Shaft, 52100 Alloy Steel, 3/8” Diameter, 8” Long, McMaster Carr), which were secured to H-supports at both ends using 16mm M4 screws (GM Lab). The top acrylic sheet was press fitted onto the H-supports to complete the primary framework. Motors for motion in the *z*-axis (two motors), *y*-axis, and *x*-axis were purchased from Digikey and assembled into their respective compartments (Figure 4).

Integration of robot motion with pneumatic regulator control: To control the robot, an Arduino Mega (Monza, IT) was utilized as the primary controller. *X* and *y* motors were controled utilizing an Adafruit motor shield (v2, Adafruit Industries) with an external 12V 2A power supply. Two A4998 stepper motor drivers (Pololu Corporation, Las Vegas, Nevada) with a 1/4 microstep were used to control both *z* stepper motors, with an addition 12V 3A power supply. To serially communicate with the five 50 psi PCD-series pressure regulators and one 100 psi PCD-series pressure regulators as shown (Movie S1). A simplex architecture operating with an inverted transistor-transistor logic protocol at a baud rate of 19200 was employed, where the gauge pressure of the devices was read via analog pins on the Arduino Mega. Communication was performed through a 8-pin mini-DIN connector and 8-pin mini-DIN to breadboard adapter.

### 0.5 Development of a graphical user interface

An ESP32-C3 M5 stamp (Digikey) was used to generate a wifi router hosting a website developed in CSS, HTML, and Javascript. This stamp was connected to the primary Arduino Mega using an serial peripheral interface communication protocol.

### 0.6 Mixing characterization of LIBRIS chip

Solutions of 10 kDa FITC-dextran (MilliporeSigma, Burlington, MA), 10 kDa rhodamine B isothiocyanate-dextran (MilliporeSigma), and 10kDa Cascade Blue®-dextran were dissolved to 10µM in ultrapure (UP) water, and filtered through 0.22µm polyethersulfone or polyvinylidene fluoride filters (ThermoFisher). Each solution served as a separate fluidic input for each of the three ethanol or aqueous phase inputs, FITC-dextran being pressurized through *i*_1_ and *i*_4_, rhodamine B-dextran through *i*_2_ and *i*_5_, and Cascade Blue®-dextran through *i*_3_ and *i*_6_. The other three fluidic inputs (either aqueous or ethanol) comprised UP water. The image of the full chip was taken where all inputs *i*_1_ and *i*_4_ carried FITC-dextran, *i*_2_ and *i*_5_ carried rhodamine B-dextran, and *i*_3_ and *i*_6_ carried Cascade Blue-dextran (Figure 2B). Each input was pressurized at the pressures used to generate the LNP libraries (Table S1), resulting in a total flow rate of 300µL/min for the ethanol phase, and 900µL/min for the aqueous phase, thus 1.2mL/min through each combined mixing unit.

Fluorescence micrographs of the entire chip was taken using a fluorescence microscope (Leica, Wetzlar, Germany). Intensity profiles were measured for all three fluorescence channels at the same location on the chip using image analysis tools in Python.

Mixing efficiency was calculated using the equation described in Section 2.2. Solutions varied from *H* = 0 (completely unmixed, prior to any mixing cycles) to *H* = 1 (completely mixed). Above *H* = 0.8, or 1− *H* = 0.2, solutions were considered completely mixed,38 validated by the fitting of an exponential decay model to each data set. Standard errors were calculated from the model fit at 10 mixing cycles for the ethanol or aqueous SHMs or 15 mixing cycles for the LNP generating SHM.

Measurements of the relative flow rates in each channel were determined by analysis of the total intensities of each dye at the fully mixed stage (1 − *H <* 0.2, after 10 mixing cycles) of a channel when normalized to the intensity of the dye at an entirely unmixed stage (before 0 mixing cycles). Thus, for *i*_1_ in *m*_1_, the relative intensity, *I*_1,1_, thus the normalized relative flow rate *Φ*_1,1_ is calculated: 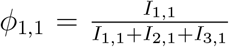 (4) where each *I*_1,1_ is calculated from: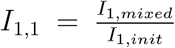 for a given fluorescence channel measurement from a given channel. To generate the images seen in Figure 2B, images were stitched together and blended using Adobe Photoshop (San Jose, CA). All mixing analysis was performed from raw images.

LNP Formulation and Characterization: LNPs were formulated using the LIBRIS chip at the designated compositional ratios (Table S1-3). For all three libraries (MC3, SM-102, C12-200), the IL and cholesterol were pooled in fluidic input *i*_1_ and the phospholipid and PEG lipid were pooled together in fluidic input *i*_3_. The stock solutions used to make the library comprised the lipids D-Lin-MC3-DMA (MedChemExpress, Monmouth Junction, NJ), 1,2-distearoyl-sn-glycero-3-phosphocholine (DSPC, Avanti Polar Lipids, Alabaster, AL), cholesterol (Sigma-Aldrich, St. Louis, MO), and DMG-PEG 2000 (Avanti Polar Lipids) at concentrations of 2.22mg/mL, 0.55mg/mL, 1.03mg/mL, and 0.26mg/mL respectively. The SM-102 library comprised SM-102 (Cayman Chemical, Ann Arbor, MI), DSPC, cholesterol, and DMG-PEG 2000 at concentrations of 2.22mg/mL, 0.49mg/mL, 0.93mg/mL, and 0.24mg/mL respectively. The C12-200 library comprised the lipids C12-200 (Cayman Chemical), 1,2-dioleoyl-sn-glycero-3-phosphoethanolamine (DOPE, Avanti Polar Lipids), cholesterol, and 1,2-distearoyl-sn-glycero-3-phosphoethanolamine-N-[amino(polyethylene glycol)-2000] (DSPE-PEG 2000, Avanti Polar Lipids) at concentrations of 2.22mg/mL, 0.66mg/mL, 1.00mg/mL, 0.38 mg/mL respectively. Input *i*_2_ comprised a solution of filtered ethanol.

The aqueous inputs *i*_4_, *i*_5_, and *i*_6_ comprised nucleic acid cargo, and two buffer inputs of 10mM citrate buffer at pH ∼ 3 (Alpha Teknova, Inc., Hollister, CA) respectively. Measures of chip repeatability and scalability were conducted using a ssDNA cargo of the sequence 5’– ATG GTT CTA GCT T – 3’ (IDT, Coralville, IA) as the nucleic acid cargo with the SM-102 lipid set. Otherwise, the nucleic acid cargo consisted of either firefly luciferase or mCherry CleanCap® mCherry mRNA (5moU) (TriLink Biotechnologies,

San Diego, CA). The LIBRIS chip was primed for 10s using UP water, then either ethanol or citrate depending on the fluidic input, each for 30s. After priming, lipid stocks were loaded, then the microcontroller was initiated, pressurizing each input to the the pressures referenced (Table S1-3). Lines were flushed and flow equilibrated for 20s, the first sample was collected for 20s, then each set of LNPs was equilibrated between for 2.6s. After sample collection, the chip was flushed with 1% Alconox (Alconox Inc., Glenn Plains, NY) for 2min, then flushed with UP water for an additional 2min.

Each output was collected in a 0.3mL 20k MWCO Pierce Microdialysis Cassette (A50472, ThermoFisher Scientific, Waltham, MA), sitting in a well of 1.2mL of 1X phosphate buffered saline (PBS, ThermoFisher Scientific). The cassettes were arranged in a 4 × 2 pattern matching the chip output setup (Figure S10-11). After collection of all 32 outputs for a given set of fluidic inputs, the dialysis cassettes were transferred to new wells of 1.2mL of PBS two additional times at intervals of 20 minutes. Samples were transferred from the cassettes and stored at 4°C.

LNPs were diluted 1:30 in 1x PBS, and, using a DynaPro Plate Reader III (Wyatt Technology, Santa Barbara, CA), dynamic light scattering was performed to quantify the hydrodynamic diameter (referred to as “size” throughout) and polydispersity index (PDI). All sizes reported are the intensity-weighted mean hydrodynamic diameters (z-averages). Standard deviation is calculated based on measurement replicates. Relative encapsulation efficiency and RNA concentration were measured by reading out the results of a RiboGreen Quant-it RNA assay kit (ThermoFisher) on an Infinite M Plex plate reader (Tecan, Männedorf, Switzerland).

### 0.7 *In vitro* characterization of mRNA transfection

HepG2 cells (HB-8065, ATCC, Manassas, VA) were cultured in DMEM (ThermoFisher) supplemented with 10% v/v fetal bovine serum (FBS, ThermoFisher) and 1% v/v Penicillin-Streptomycin (ThermoFisher). Cells were plated at 40k cells/100µL of media in a 96-well plate, then dosed after 24hr with either mCherry LNPs from the MC3, SM-102, or C12-200 libraries at a dose of 10ng mRNA/10k cells. After another 24hr, cells underwent preparation for flow cytometry as follows.

Each well of cells was washed with 100µL of 1× PBS, then treated with 60µL of 0.25% Trypsin-EDTA (ThermoFisher) for 6 min at 37°C. Each well was diluted in 100µL of DMEM, then spun down at 0.3rcf for 5min.

LIVE/DEAD™ Fixable Green Dead Cell Stain (ThermoFisher) was diluted in 1x PBS to a concentration of 1µL of dye per 5mL of PBS. Media was aspirated, and 50µL of dye was added to each well. Each well was agitated using a multichannel pipette and left to rest covered from light for 30min at 25°C. 100µL of 1x TE buffer (ThermoFisher) was added to each well, and the plate was centrifuged at 0.3rcf for 5 min. TE was aspirated, and 70µL of 2% v/v neutral buffered formalin (NBF, ThermoFisher) diluted in 1x PBS was added to each well, left to incubate covered from light for 10min, then the plate was centrifuged at 0.6rcf for 5min. 150µL of 1x TE buffer was added to the cells, the cells were centrifuged again at 0.6rcf for 5min, then resuspended in 150µL of 1x PBS and stored at 4°C until run on the cytometer.

A Guava® easyCyte™ HT System (Luminex, Austin, TX) was used to perform the cytometry experiment. Samples were gated according to the following scheme (Figure S18).

### 0.8 *In vivo* characterization

Candidates were selected from the *in vitro* assay for high performance (C12-200 F4), for poor performance (C12-200 F24), and from the previously optimized standard formulation for C12-200 [6] for validation *in vivo*. Compositional ratios for each particle were calculated from the relative flow rates and input concentrations of F4 and F24 collected from the LIBRIS chip as they corresponded to their *in vitro* performance. Ethanol stocks were mixed where all of the lipids were pooled at the calculated compositional ratios (Table S1-3), and were precipitated together with 5-methoxyuridine CleanCap® Firefly Luciferase mRNA (Trilink) dissolved in pH∼ 3 10mM citrate buffer (ThermoFisher) through a single microfluidic mixing device as described in previous literature [20, 44]. The relative flow rates of the ethanol to aqueous phases were 1:3 respectively, with a total flow rate of 1.2mL as dictated by displacement driven flow using a syringe pump (Pump 33 DDS, Harvard Apparatus, Holliston, MA). Samples were collected in 20-kDa MWCO Slide-A-Lyzer™ dialysis cassettes (ThermoFisher) and dialyzed against 1x PBS (ThermoFisher) for 1.5hr before continuing to analysis.

Female C57BL/6 mice between six and eight weeks old (Jackson Labs, Bar Harbor, ME) were used for all animal experimentation, performed in approval of the University of Pennsylvania Institutional Animal Care and Use Committee (IACUC, protocol number 806540).

The temperature of the animal housing facility was 22±2 °C, held at a 12hour dark/light cycle, with 40-70% air humidity. Randomly allocated mice for each group were injected intravenously (i.v.) at a dose of 0.1mg mRNA/kg (2µg per mouse, around 100µL of volume through a tail vein injection). After 6hr, mice were injected intraperitonealy (i.p.) with 200µL of D-luciferin potassium salt (ThermoFisher) at a concentration of 15mg/mL. Mice were sacrificed 10min post i.p. injection, and their hearts, lungs, kidneys, livers, and spleens were dissected and imaged using a Lumina S3 in vivo imaging system (IVIS, PerkinElmer, Waltham, MA). Bioluminescence analysis was performed in Living Image 4.7.3 software (PerkinElmer).

Statistical analysis Statistical analyses and plotting were performed in Python using the scikit-learn or SciPy packages or in GraphPad Prism 10 with significance level *α* = 0.05. Data are represented as mean *±* s.d. Oneway ANOVA followed by Tukey’s HSD test or the LSD test followed by a Bonferroni correction were performed for multiple comparisons between groups.

## Acknowledgements

M.J.M. acknowledges support from a US National Institutes of Health Director’s New Innovator Award (DP2 TR002776), a Burroughs Wellcome Fund Career Award at the Scientific Interface (CASI), a US National Science Foundation CAREER award (CBET-2145491), an American Cancer Society Research Scholar Grant (RSG-22-122-01-ET), and the Cystic Fibrosis Foundation (MITCHE24I0). D.A.I. acknowledges support from the Wellcome Leap R3 program, NSF MRSEC Grant (DMR-2309034), NSF Biofoundry, and the Center for Precision Engineering for Health at University of Pennsylvania. A.R.H., S.J.S., I.B.N., and H.M.Y. were supported by US National Science Foundation Graduate Research Fellowships. M.S.P. was supported by the National Institute of Dental and Craniofacial Research of the US National Institutes of Health (T90DE030854). Chip microfabrication was performed in the Quattrone Nanofabrication Facility at the Singh Center for Nanotechnology at the University of Pennsylvania. Parts were taken from the GM Lab at the University of Pennsylvania.

## Author Contributions

A.R.H., S.J.S., M.J.M., and D.A.I. conceived and designed the experiments. A.R.H. wrote the manuscript and prepared the figures. A.R.H., G.A.D., A.S.R., I.B.N., M.S.P., S.Z., N.Y.M., and H.M.Y. performed the experiments. A.R.H. and G.A.D. analyzed the data. G.A.D, I.B.N. and J.R.B. assisted in design of the plate robotic interface. N.S. advised parts of the chip fabrication. A.R.H., M.J.M., and D.A.I. reviewed and edited the manuscript. M.J.M. and D.A.I. supervised the entire project. All authors approved the final manuscript for submission.

## Data Availability

All data relevant to this study are included in the article and its Supplementary Information or are available from A.R.H., M.J.M., or D.A.I. upon reasonable request. Raw images are available from A.R.H., M.J.M., or D.A.I. upon reasonable request.

