## Supplementary Information for "Automated and parallelized microfluidic generation of large and precisely-defined lipid nanoparticle libraries"

May 1, 2025

Andrew R. Hanna, Sarah J. Shepherd, Gregory A. Datto, Isabel B. Navarro, Adele S. Ricciardi, Marshall S. Padilla, Neha Srikumar, Shuran Zhang, Hannah M. Yamagata, Nova Y. Meng, Joshua R. Buser, Michael J. Mitchell, David Issadore

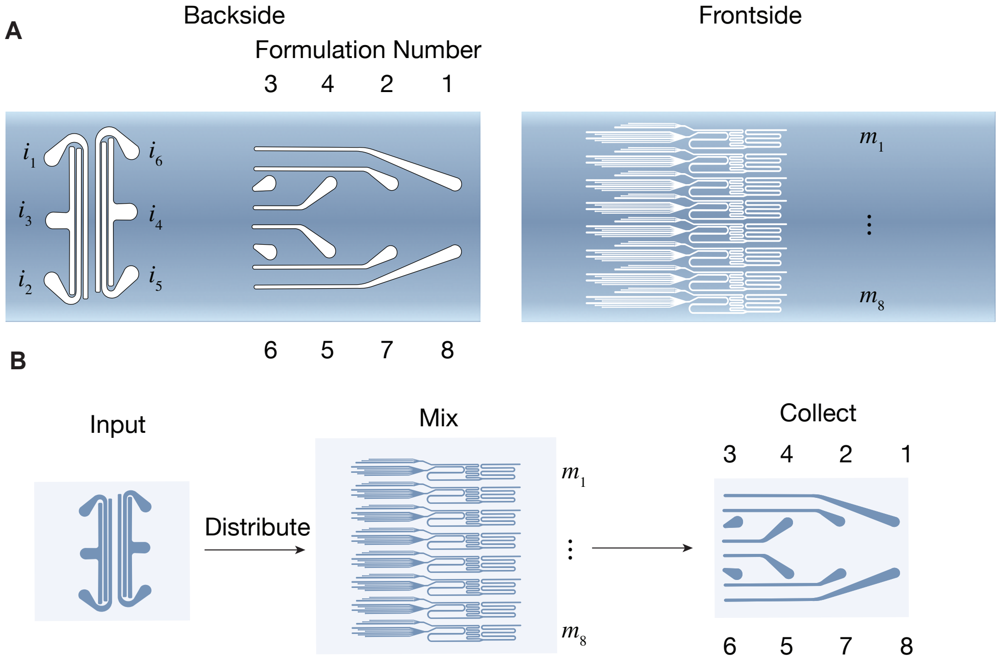


Figure 1: (A) Schematic overview of outputs and inputs of chip as referred to in manuscript. Formulations 1 – 8 collected from outputs as labelled, each corresponding to an LNP generator 1 – 8 as seen on the frontside schematic. (B) Overview of pathway of fluids through chip.

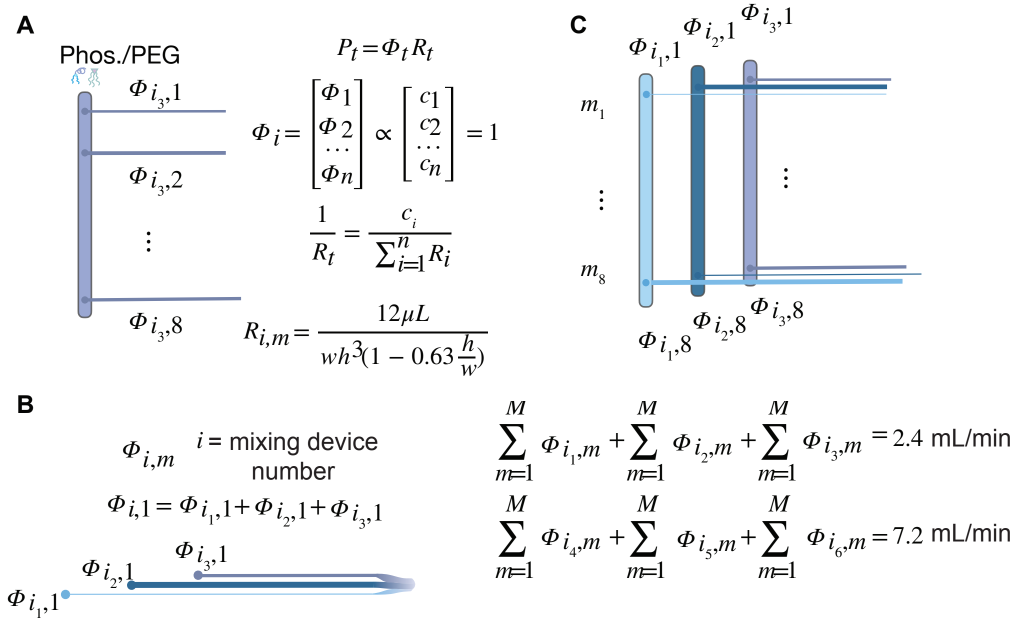


Figure 2: Derivation of design rules for fluidic distribution across resistors for a given input. (A) Distribution of fluids across all mixing units through a single input is governed by the desired composition matrix shown, where all possible compositional flow rates are normalized to the total flow rate through a single delivery channel. The necessary fluidic resistances are calculated by the relation $\frac{P_i}{\Phi_i} = \left(\sum_{m=1}^{M} \frac{1}{R_{i,m}}\right)^{-1}$ (B) The total flow rate through single mixing unit is the sum of the flow rates through each of the three resistor $R_{i,m}$ where $i = 1 - 3$ for the ethanol inputs or $4 - 6$ for the aqueous inputs. The sum of these multiple flow rates through a single LNP generator is held at either 300µL/min for the ethanol phase and 900µL/min for the aqueous phase. The variation in the flow rates occurs where one input is held fixed at a given set of pressures, and the other two inputs vary inversely with one another. (C) The flow rates are distributed as if through a network of parallel resistors. To prevent backflow between resistors, the resistances of all SHMs for each LNP generator are at least 5× lower than the the resistance of any resistor $R_{i,m}$ (Figure 2B).

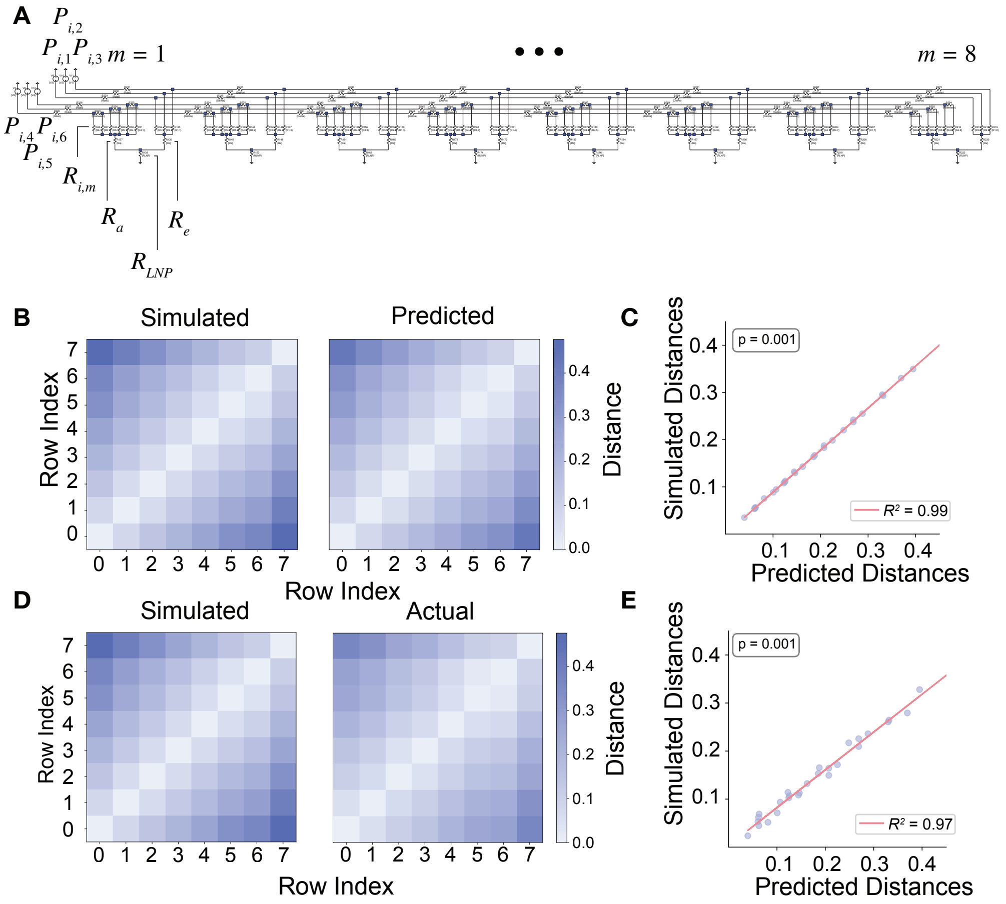


Figure 3: (A) LTSpice model of circuit, where blue boxes represent junctions. Pressure (Pa) is analogized as voltage, where each pressure source is labelled as the pressure at the inlet number i for a given pressure setting n (Pi,n). Flow rate ($\frac{mm^3}{s}$) is analogized to current. Hydraulic or fluidic resistance ($\frac{Pa \times s}{mm^3}$) is analogized to electrical resistance. Pairwise distance analyses were performed between relative flow rate values derived from the LTSpice simulation and both (B, C) predicted values ($R^2 = 0.99$) and (D, E) actual values ($R^2 = 0.97$).

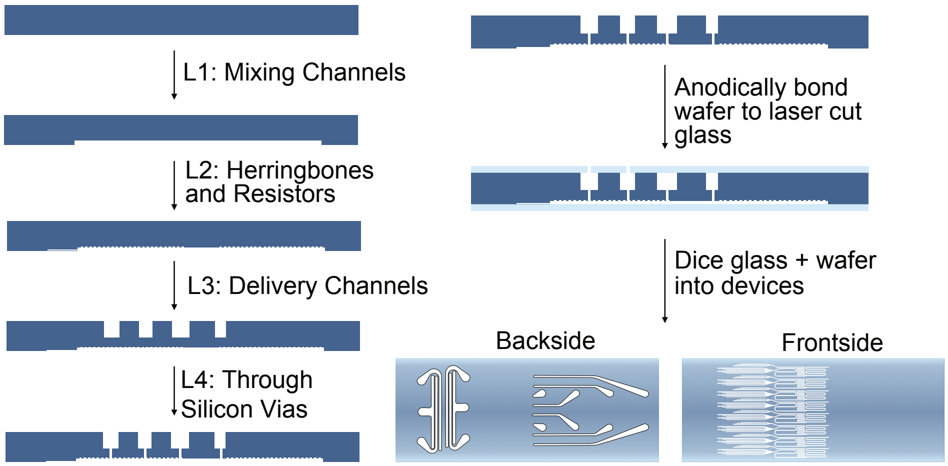


Figure 4: Overall fabrication overview for the device. Layer depths are as follows: L1: 85µm, L2: 20µm, L3: 365µm, L4: ≈120µm. After patterning, the wafer is anodically bonded to Borofloat 33 glass on both sides then diced into single dies.

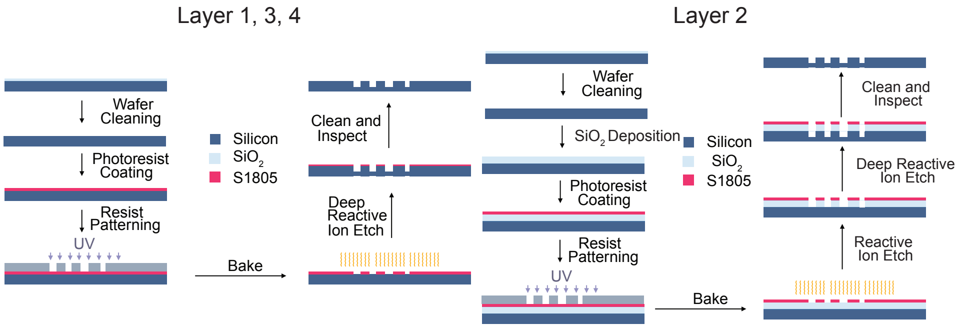


Figure 5: Schematic fabrication overview for a single layer. Layers 1, 3, and 4 comprise the same design strategy, while Layer 2 for the herringbones applies a 500nm hard mask to prevent undesired etching of the corners of the mixing channels.

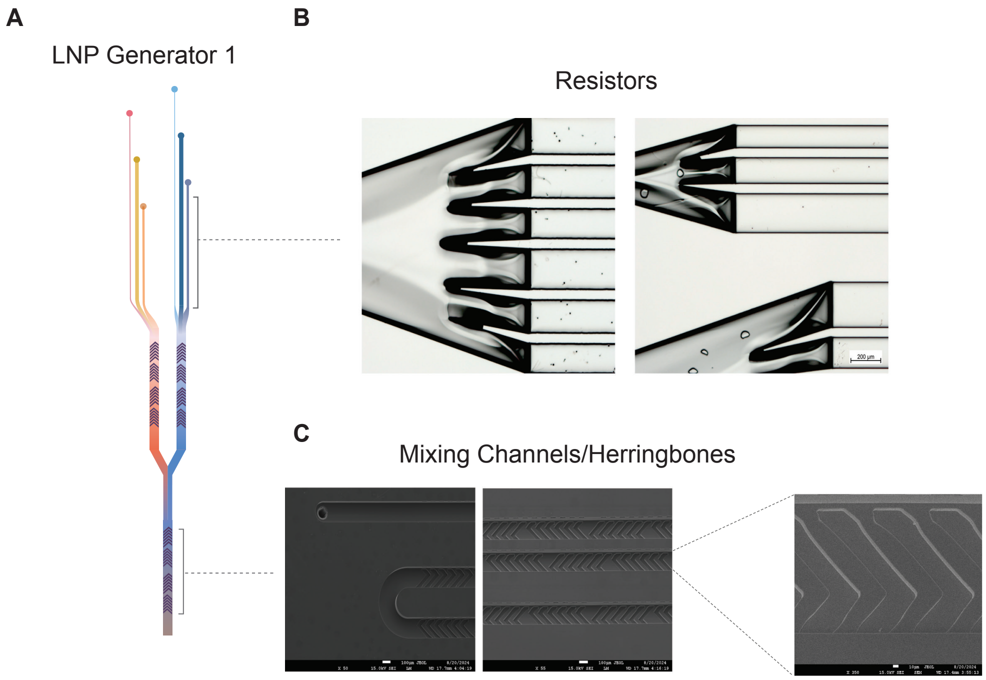


Figure 6: (A) Schematic of single mixing device. (B) Images of resistor and entrance to (right) aqueous mixing unit and (left) ethanol mixing unit taken with light microscope (Scale Bar: 200µm). (C) Scanning electron micrograph of herringbones, mixing channel (Scale Bar: 100µm), and close up of herringbones (Scale Bar: 10µm).

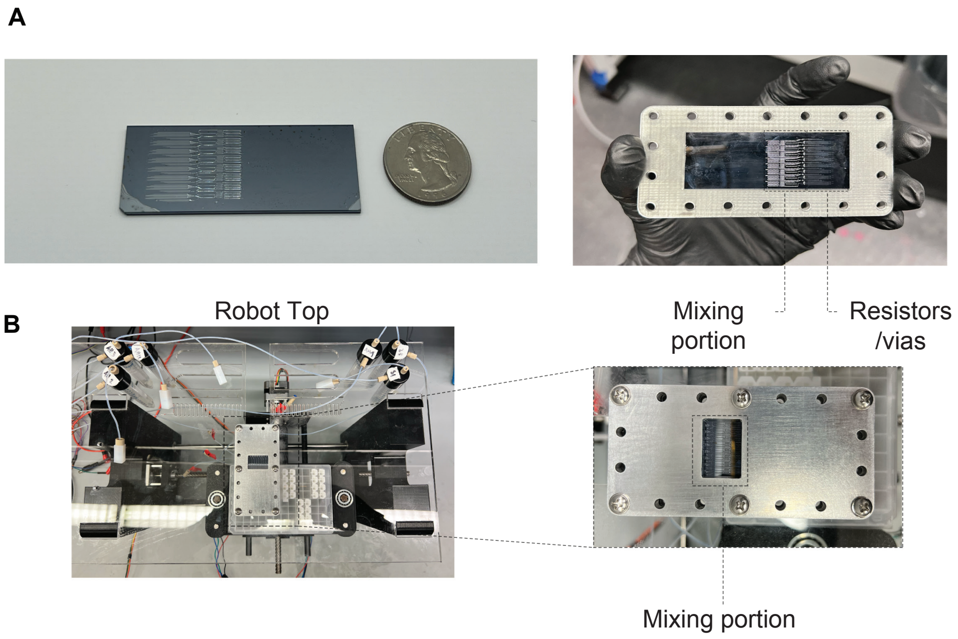


Figure 7: Images of the single chip die (A) next to a US quarter for scale, (B) surrounded by an acrylic alignment piece (¡1.5mm thick) to align the device to the center of the aluminum holder pieces using exterior bolts, and (C, D) positioned and fastened to the top plate of the plate robot within its holder. Each of the aluminum pieces is 6.4mm thick while the device itself is 1.5mm thick. Tubing is attached to the device from the bottom side of the aluminum holder using a set of $\frac{1}{4}$"-28 threaded ports and fittings holding tubing fastened by flangeless ferrules.

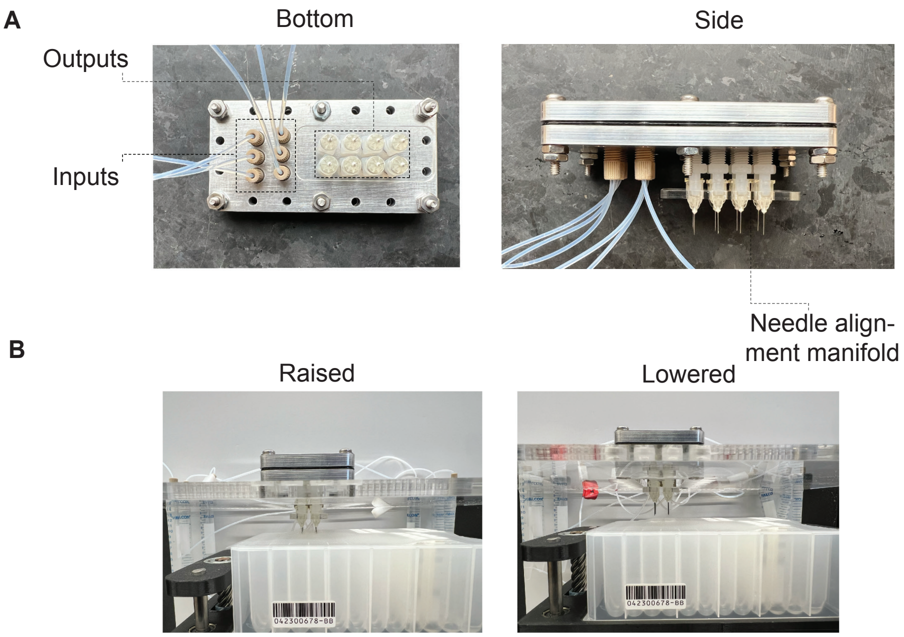


Figure 8: Labeled diagram of the input tubing, output needles, and needle manifold. (A) Fluid enters each of the respective inputs, mapped in Figure S1. Needles are aligned and additionally secured using a laser-cut acrylic manifold. (B) Images of needles puncturing and aligning with dialysis cassettes in plate beneath.

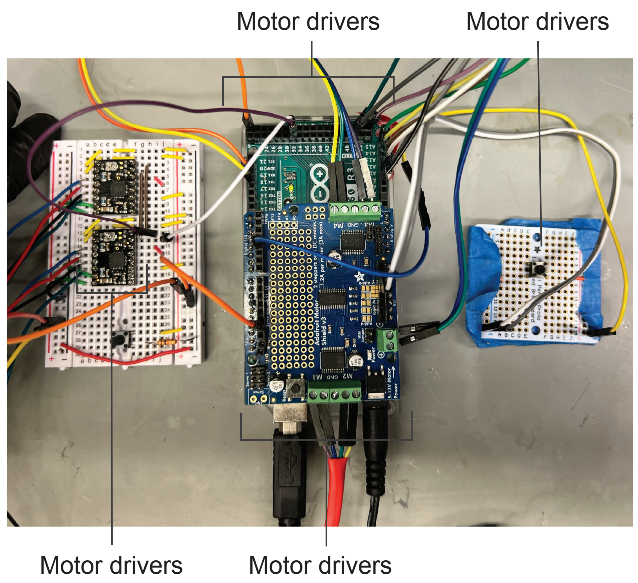


Figure 9: Image of the microcontroller and circuitry for controlling motors. The Arduino Mega connects to separate boards which drive the $x$, $y$, and $z$ motors (board not shown). Initiation is driven by the control button. Serial communication with the pressure regulators is allowed by a separate series of circuits (not shown) which connects a series of control pins from an 8-pin mini-DIN connector to control via the Arduino.

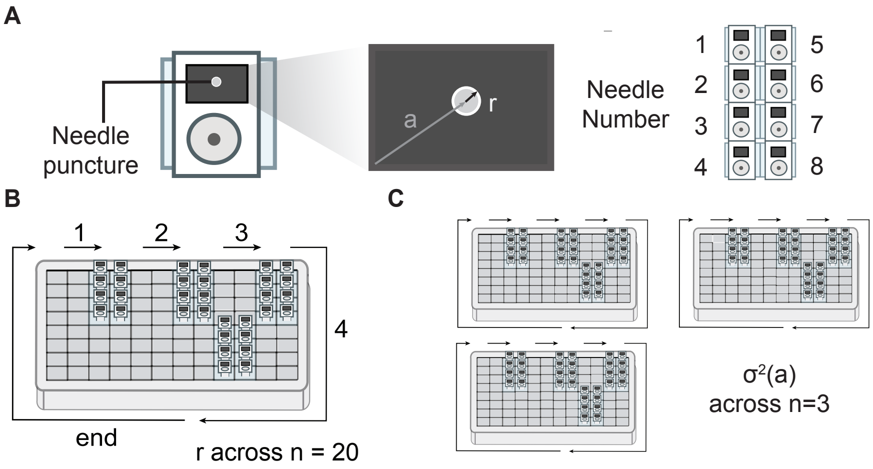


Figure 10: Schematics of plate robot consistency experiments. (A) Absolute distance measured between plates was measured as the Euclidean distance a from the bottom right corner of the cassette, and the radius $r$ of puncture marks is measured as shown to measure the variability in motion across collection positions on a single plate. Needle numbers in bar plots are measured as shown (right). (B) Each of the collection positions on the plate is shown by a $4 \times 2$ arrangement of dialysis cassettes, followed by a waste well. Figure 4F shows data collected across these positions 20 times. The process was repeated across (C) three separate plates, and the variance of collection at each of the four positions ($n = 3$) was calculated and compared between needle positions. The averages of the absolute position a for each of the needle positions was also pooled for each of the plates and compared.

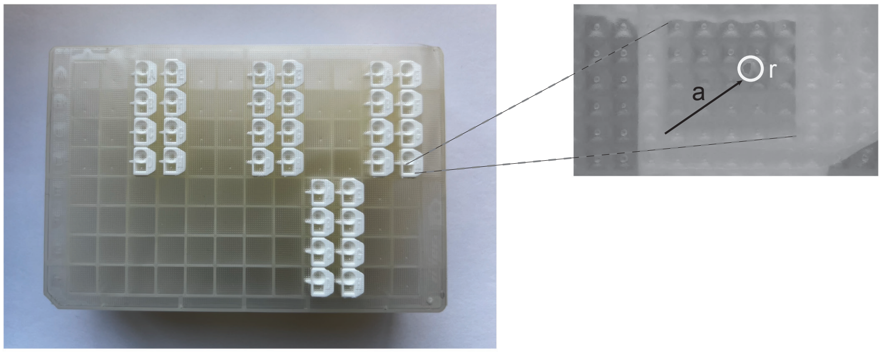


Figure 11: The puncture position was measured from the corner of these dialysis cassettes as shown. Images were taken under the microscope which was used as a part of the robot monitoring system and analyzed using FIJI ImageJ, fitting an ellipse to the puncture marks and calculating the radius circumscribed by the major axis.

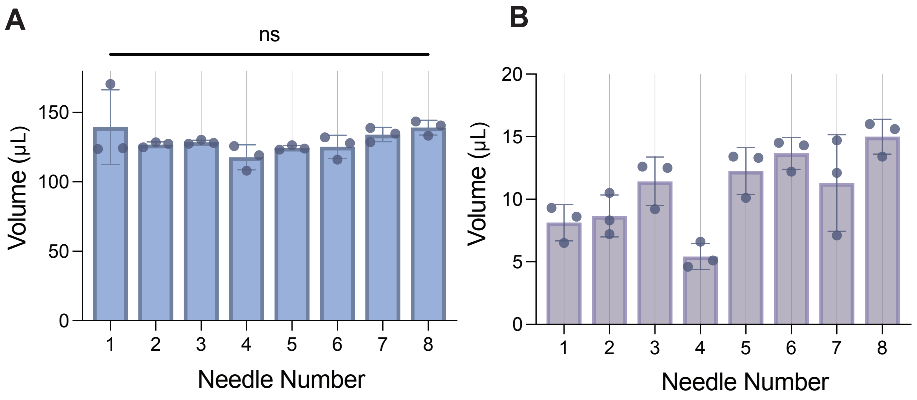


Figure 12: Measurements of consistency and minimum volume collected by the LIBRIS platform. (A) Sample volume was collected for 10s at $n = 3$ collection positions at the second operating pressure setting (Table S1), and sample volume was collected in sets of prehydrated cassettes in a plate. The efficiency of volume collection after the entirety of processing (collection, removal from cassettes, measurement of volume) was between 60-70% for each sample. (B) Sample volume was collected for 1s and measured. Collection efficiency through the entirety of processing was on average about 5%. The lower efficiency is likely from the larger relative proportion of volume clinging to interfaces (cassette walls, pipette tip walls, tube walls) through the collection process.

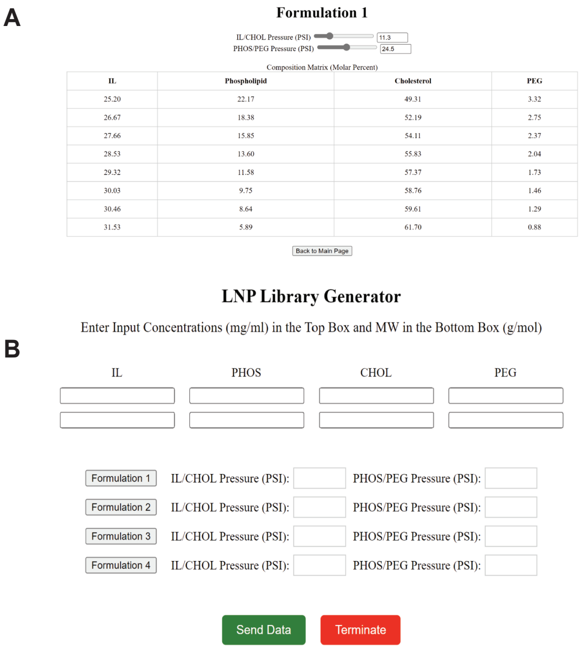


Figure 13: Screenshots of the graphical user interface for use of the LIBRIS platform. (A) Users can input their starting concentrations and proceed to the following screen (B) where they are shown a sliding scale, demonstrating the range of compositions produced at a given set of inversely correlated pressure settings. After the user has selected their desired pressure settings based on the molar ratios which they desire, they can enter those pressures into the numbered formulation entry settings in screen (A) and wirelessly communicate with the microcontroller, then using the button initiatior (Figure S9) to begin the run, collecting their desired formulations.

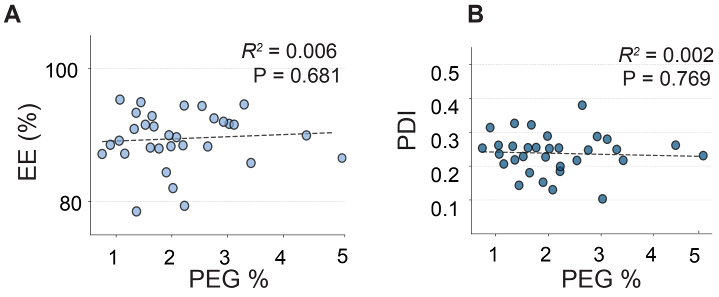


Figure 14: Unlike size (nm), neither (E) EE% ($R^2$ = 0.006) nor (F) PDI ($R^2$ = 0.002)independently demonstrates Pearson correlation with the molar percentage of PEG lipid.

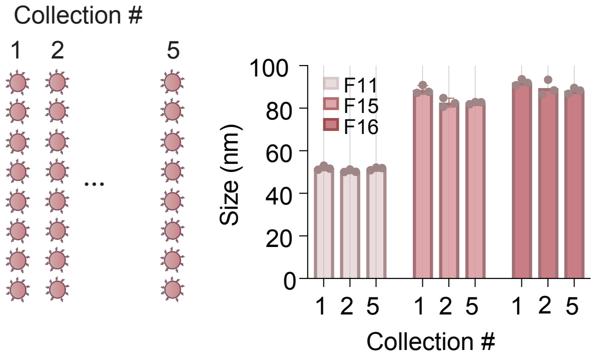


Figure 15: Particle sizes remain constant after 5 separate collections using the same operating parameters, indicating that the chip operation does not contribute to any differences in particle physicochemical characteristics over the course of the experiment.

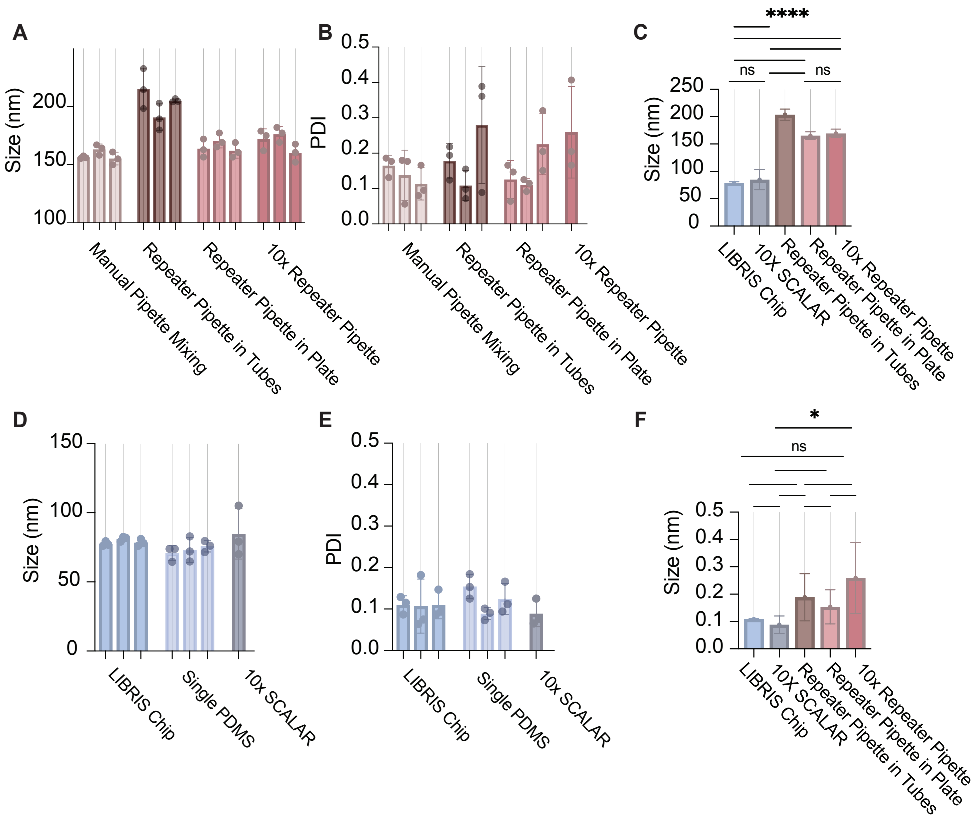


Figure 16: (A-B) Using a manual pipette, an automated pipette in two unique output geometries, we mixed 50µL of SM-102 lipids (Methods) dissolved in EtOH with 150µL of ssDNA nucleic acid cargo dissolved in 10mM Citrate Buffer at pH 3, at the maximum speed possible for mixing. (D-E) Using the same proportion of lipids and RNA, we generated LNPs using our LIBRIS chip, a control single PDMS device [1], and a 10× parallelized version of a SHM device [2]. We demonstrate that LNPs generated using the LIBRIS chip compare in physicochemical properties with the 10× SCALAR chip (C, F), and are significantly smaller than those made using pipette mixing methods. Thus, we show that according to physicochemical properties, the LIBRIS chip can produce particles at discovery scale that can be exactly reproduced at any scale. Further *in vitro* and *in vivo* validation of this point can be observed here [2]. (C, Size: $W = 710.7$, **** $P < 0.0001$; LIBRIS vs. 10× SCALAR: $P = 0.9969$ by Welch's ANOVA and Dunnett's post-hoc test, $n = 9$. F, PDI: $W = 5.292$, * $P = 0.0384$; LIBRIS vs. 10× SCALAR: $P = 0.9003$ by Welch's ANOVA and Dunnett's post-hoc test, $n = 9$.

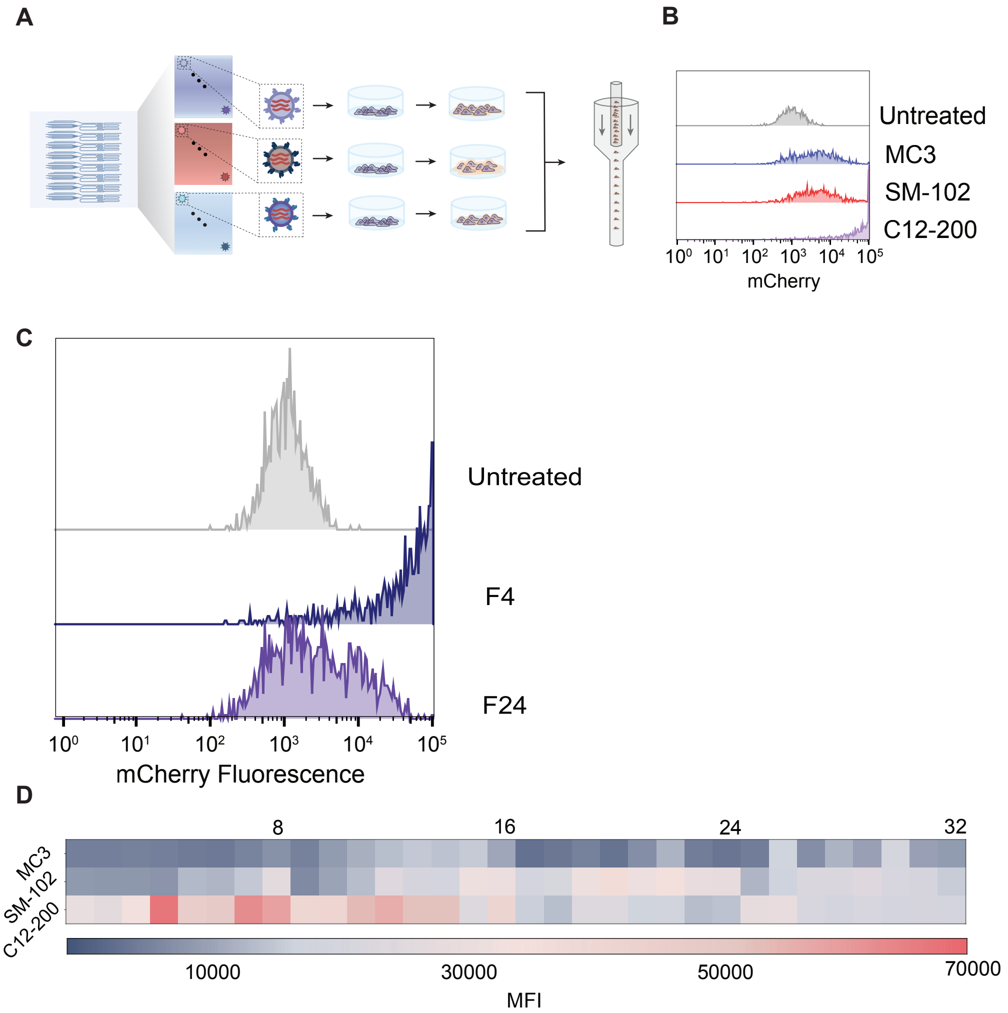


Figure 17: (A) Schematic of LNP generation via LIBRIS chip and testing using flow cytometry. (B) Representative histogram from flow cytometry of samples selected from each of the compositional groups screened. (C) Flow cytometry histogram for HepG2 cells treated by formulations further assayed in the in vivo screen. LNPs generated mRNA encoding for mCherry were generated using the LIBRIS chip and dosed at 40ng of mRNA/40,000 cells. Shifts in fluorescence intensity of the samples collected demonstrate increases in fluorescence intensity relative to untreated samples. (D) Mean fluoresence intensity (MFI) of all 96 samples across library.

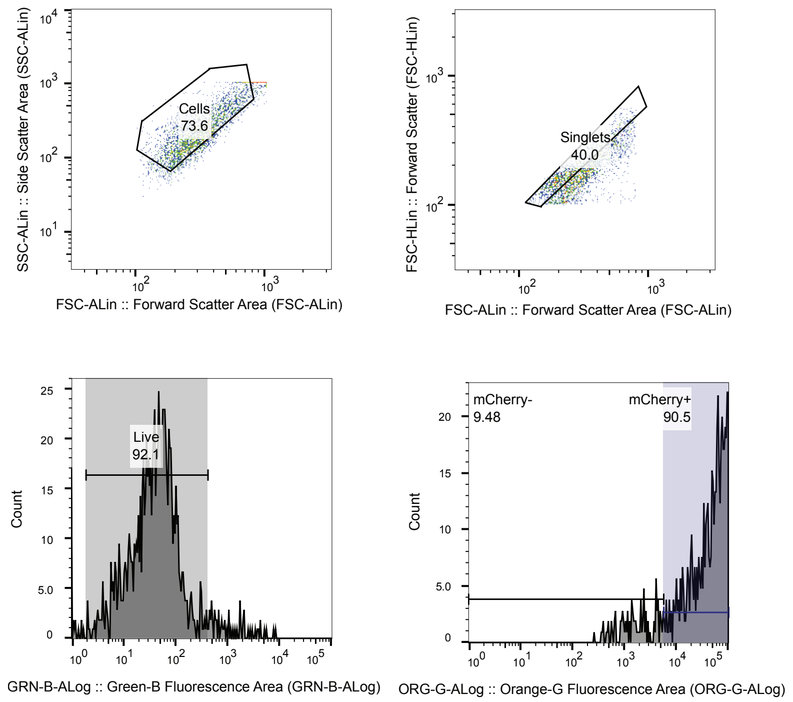


Figure 18: Representative flow cytometry gating scheme for evaluatign the proportion of transfected (mCherry$^{+}$) HepG2 cells.

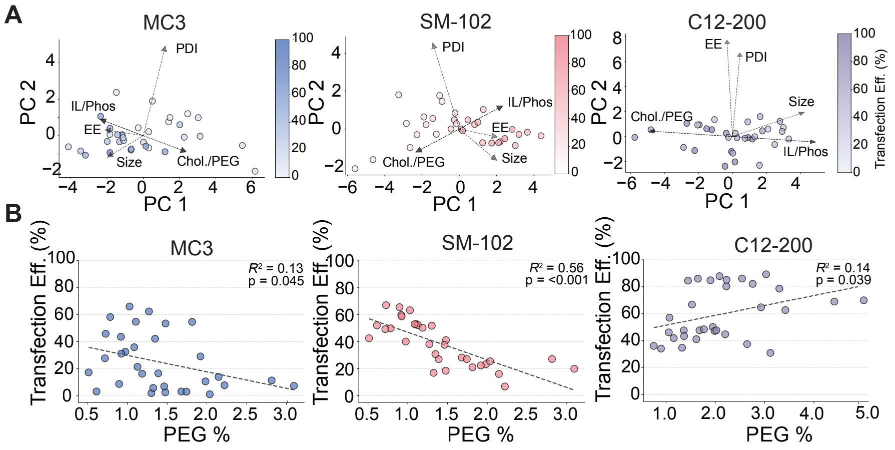


Figure 19: (A) Principal component analyses of compositional and physicochemical variables for each library analyzed against transfection efficiency. Loadings are as follows: MC3: IL, PEG, Chol, PEG (0.42, 0.16); EE (0.36, 0.05); Size (0.34, 0.21); PDI (0.23, 0.93); SM-102: IL, PEG, Chol, PEG (0.41, 0.22); EE (0.35, 0.07); Size (0.36, 0.29); PDI (0.27, 0.85); C12-200: IL, PEG, Chol, PEG (0.46, 0.04); EE (0.03, 0.74); Size (0.40, 0.18); PDI (0.04, 0.64). (B) Pearson correlation analysis of % mCherry$^+$ cells and molar percentage of PEG (see insets for $R^2$ and $P$ values).

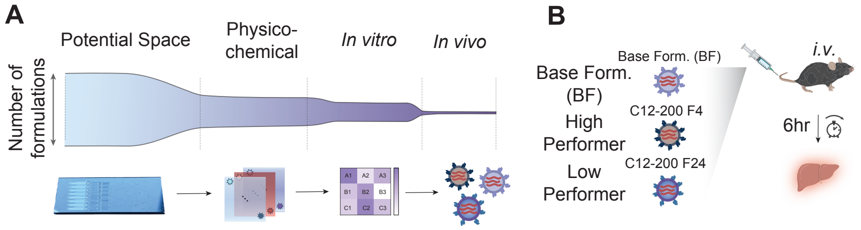


Figure 20: Schematic of narrowing the LNP design space using the LIBRIS chip.

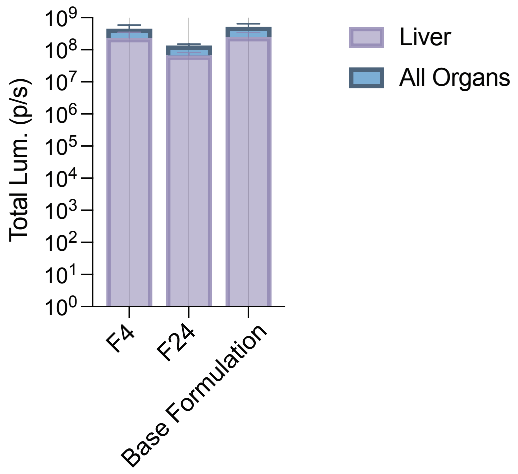


Figure 21: Quantification of luciferase signal by IVIS in liver versus all organs combined.

| Form. | Predicted | | | Simulated | | | Actual | | | Pressure (psi) | | |
|---|---|---|---|---|---|---|---|---|---|---|---|---|
| | $i_1$ | $i_2$ | $i_3$ | $i_1$ | $i_2$ | $i_3$ | $i_1$ | $i_2$ | $i_3$ | $P_1$ | $P_2$ | $P_3$ |
| 1 | 0.181 | 0.271 | 0.549 | 0.135 | 0.274 | 0.591 | 0.214 | 0.251 | 0.535 | 16 | 24 | 24 |
| 2 | 0.205 | 0.307 | 0.488 | 0.156 | 0.318 | 0.526 | 0.226 | 0.292 | 0.483 | | | |
| 3 | 0.223 | 0.335 | 0.442 | 0.172 | 0.352 | 0.476 | 0.284 | 0.312 | 0.404 | | | |
| 4 | 0.241 | 0.362 | 0.397 | 0.188 | 0.385 | 0.426 | 0.305 | 0.306 | 0.388 | | | |
| 5 | 0.259 | 0.388 | 0.353 | 0.204 | 0.418 | 0.377 | 0.315 | 0.363 | 0.322 | | | |
| 6 | 0.276 | 0.414 | 0.310 | 0.220 | 0.451 | 0.329 | 0.328 | 0.388 | 0.284 | | | |
| 7 | 0.287 | 0.431 | 0.282 | 0.228 | 0.474 | 0.298 | 0.338 | 0.391 | 0.271 | | | |
| 8 | 0.318 | 0.477 | 0.205 | 0.256 | 0.530 | 0.214 | 0.353 | 0.461 | 0.186 | | | |
| 9 | 0.226 | 0.226 | 0.549 | 0.205 | 0.205 | 0.591 | 0.326 | 0.187 | 0.487 | 20 | 20 | 24 |
| 10 | 0.256 | 0.256 | 0.488 | 0.237 | 0.237 | 0.526 | 0.367 | 0.186 | 0.447 | | | |
| 11 | 0.279 | 0.279 | 0.442 | 0.262 | 0.262 | 0.476 | 0.368 | 0.217 | 0.415 | | | |
| 12 | 0.301 | 0.301 | 0.397 | 0.287 | 0.287 | 0.426 | 0.416 | 0.222 | 0.362 | | | |
| 13 | 0.323 | 0.323 | 0.353 | 0.311 | 0.311 | 0.377 | 0.442 | 0.239 | 0.319 | | | |
| 14 | 0.345 | 0.345 | 0.310 | 0.335 | 0.335 | 0.329 | 0.457 | 0.264 | 0.279 | | | |
| 15 | 0.359 | 0.359 | 0.282 | 0.351 | 0.351 | 0.298 | 0.464 | 0.277 | 0.260 | | | |
| 16 | 0.397 | 0.397 | 0.205 | 0.393 | 0.393 | 0.214 | 0.502 | 0.308 | 0.190 | | | |
| 17 | 0.271 | 0.181 | 0.549 | 0.274 | 0.135 | 0.591 | 0.385 | 0.106 | 0.509 | 24 | 16 | 24 |
| 18 | 0.307 | 0.205 | 0.488 | 0.318 | 0.156 | 0.526 | 0.428 | 0.114 | 0.458 | | | |
| 19 | 0.335 | 0.223 | 0.442 | 0.352 | 0.172 | 0.476 | 0.462 | 0.142 | 0.396 | | | |
| 20 | 0.362 | 0.241 | 0.397 | 0.385 | 0.188 | 0.426 | 0.492 | 0.143 | 0.365 | | | |
| 21 | 0.388 | 0.259 | 0.353 | 0.418 | 0.204 | 0.377 | 0.520 | 0.155 | 0.325 | | | |
| 22 | 0.414 | 0.276 | 0.310 | 0.451 | 0.220 | 0.329 | 0.559 | 0.165 | 0.276 | | | |
| 23 | 0.431 | 0.287 | 0.282 | 0.474 | 0.228 | 0.298 | 0.552 | 0.184 | 0.264 | | | |
| 24 | 0.477 | 0.318 | 0.205 | 0.530 | 0.256 | 0.214 | 0.597 | 0.216 | 0.187 | | | |
| 25 | 0.316 | 0.135 | 0.549 | 0.343 | 0.066 | 0.591 | 0.533 | 0.063 | 0.404 | 28 | 12 | 24 |
| 26 | 0.359 | 0.154 | 0.488 | 0.399 | 0.075 | 0.526 | 0.535 | 0.052 | 0.414 | | | |
| 27 | 0.391 | 0.167 | 0.442 | 0.442 | 0.082 | 0.476 | 0.556 | 0.075 | 0.369 | | | |
| 28 | 0.422 | 0.181 | 0.397 | 0.484 | 0.090 | 0.426 | 0.585 | 0.065 | 0.350 | | | |
| 29 | 0.453 | 0.194 | 0.353 | 0.526 | 0.097 | 0.377 | 0.626 | 0.065 | 0.310 | | | |
| 30 | 0.483 | 0.207 | 0.310 | 0.567 | 0.104 | 0.329 | 0.648 | 0.085 | 0.267 | | | |
| 31 | 0.503 | 0.216 | 0.282 | 0.596 | 0.105 | 0.298 | 0.674 | 0.075 | 0.251 | | | |
| 32 | 0.556 | 0.238 | 0.205 | 0.668 | 0.119 | 0.214 | 0.722 | 0.091 | 0.187 | | | |
| 33 | 0.129 | 0.259 | 0.612 | 0.066 | 0.343 | 0.591 | 0.163 | 0.241 | 0.596 | 12 | 24 | 28 |
| 34 | 0.149 | 0.298 | 0.553 | 0.075 | 0.399 | 0.526 | 0.170 | 0.296 | 0.534 | | | |
| 35 | 0.164 | 0.329 | 0.507 | 0.082 | 0.442 | 0.476 | 0.207 | 0.329 | 0.464 | | | |
| 36 | 0.180 | 0.360 | 0.461 | 0.090 | 0.484 | 0.426 | 0.222 | 0.321 | 0.456 | | | |
| 37 | 0.195 | 0.390 | 0.414 | 0.097 | 0.526 | 0.377 | 0.220 | 0.367 | 0.413 | | | |
| 38 | 0.211 | 0.421 | 0.368 | 0.104 | 0.567 | 0.329 | 0.183 | 0.407 | 0.411 | | | |
| 39 | 0.221 | 0.442 | 0.337 | 0.105 | 0.596 | 0.298 | 0.242 | 0.408 | 0.350 | | | |
| 40 | 0.250 | 0.500 | 0.251 | 0.119 | 0.668 | 0.214 | 0.264 | 0.477 | 0.259 | | | |

Table 1: Predicted, simulated, and actual relative flow rates of the LIBRIS chip. Each pressure setting listed corresponds to $i_1 - i_3$. Pressure settings for $i_4 - i_6$ are $1.5\times$ the values of each corresponding inlet $i_1 - i_3$. Each set of pressures listed corresponds to a set of eight formulations.

| **Form.** | **MC3** | **DSPC** | **Chol.** | **PEG** |
|---|---|---|---|---|
| 1 | 42.61 | 21.36 | 32.83 | 3.20 |
| 2 | 44.20 | 18.91 | 34.05 | 2.84 |
| 3 | 47.67 | 13.57 | 36.73 | 2.03 |
| 4 | 48.47 | 12.35 | 37.34 | 1.85 |
| 5 | 49.87 | 10.19 | 38.42 | 1.53 |
| 6 | 50.76 | 8.81 | 39.11 | 1.32 |
| 7 | 51.14 | 8.22 | 39.40 | 1.23 |
| 8 | 52.86 | 5.58 | 40.72 | 0.84 |
| 9 | 47.30 | 14.14 | 36.44 | 2.12 |
| 10 | 48.76 | 11.89 | 37.57 | 1.78 |
| 11 | 49.25 | 11.13 | 37.94 | 1.67 |
| 12 | 50.75 | 8.83 | 39.10 | 1.32 |
| 13 | 51.64 | 7.45 | 39.79 | 1.12 |
| 14 | 52.33 | 6.40 | 40.31 | 0.96 |
| 15 | 52.65 | 5.91 | 40.56 | 0.89 |
| 16 | 53.84 | 4.08 | 41.47 | 0.61 |
| 17 | 48.21 | 12.74 | 37.14 | 1.91 |
| 18 | 49.58 | 10.63 | 38.19 | 1.59 |
| 19 | 50.82 | 8.72 | 39.15 | 1.31 |
| 20 | 51.51 | 7.65 | 39.69 | 1.15 |
| 21 | 52.24 | 6.53 | 40.25 | 0.98 |
| 22 | 53.08 | 5.23 | 40.90 | 0.78 |
| 23 | 53.17 | 5.10 | 40.97 | 0.76 |
| 24 | 54.27 | 3.40 | 41.81 | 0.51 |
| 25 | 51.42 | 7.80 | 39.61 | 1.17 |
| 26 | 51.32 | 7.95 | 39.54 | 1.19 |
| 27 | 52.00 | 6.91 | 40.06 | 1.04 |
| 28 | 52.41 | 6.27 | 40.38 | 0.94 |
| 29 | 53.07 | 5.26 | 40.88 | 0.79 |
| 30 | 53.61 | 4.42 | 41.30 | 0.66 |
| 31 | 53.87 | 4.02 | 41.50 | 0.60 |
| 32 | 54.65 | 2.83 | 42.10 | 0.42 |

Table 2: Molar compositions of MC3 particle library determined by the actual relative flow rates.

| Form. | SM-102 | DSPC | Chol. | PEG |
|---|---|---|---|---|
| 1 | 42.60 | 21.34 | 32.85 | 3.21 |
| 2 | 44.19 | 18.90 | 34.07 | 2.84 |
| 3 | 47.66 | 13.56 | 36.74 | 2.04 |
| 4 | 48.45 | 12.34 | 37.36 | 1.85 |
| 5 | 49.85 | 10.18 | 38.44 | 1.53 |
| 6 | 50.75 | 8.80 | 39.13 | 1.32 |
| 7 | 51.13 | 8.22 | 39.42 | 1.24 |
| 8 | 52.84 | 5.58 | 40.74 | 0.84 |
| 9 | 47.29 | 14.13 | 36.46 | 2.12 |
| 10 | 48.75 | 11.88 | 37.59 | 1.79 |
| 11 | 49.24 | 11.13 | 37.96 | 1.67 |
| 12 | 50.73 | 8.83 | 39.12 | 1.33 |
| 13 | 51.63 | 7.45 | 39.81 | 1.12 |
| 14 | 52.31 | 6.39 | 40.33 | 0.96 |
| 15 | 52.63 | 5.90 | 40.58 | 0.89 |
| 16 | 53.82 | 4.08 | 41.49 | 0.61 |
| 17 | 48.19 | 12.74 | 37.16 | 1.91 |
| 18 | 49.56 | 10.63 | 38.21 | 1.60 |
| 19 | 50.81 | 8.71 | 39.17 | 1.31 |
| 20 | 51.50 | 7.65 | 39.71 | 1.15 |
| 21 | 52.22 | 6.53 | 40.27 | 0.98 |
| 22 | 53.07 | 5.23 | 40.92 | 0.79 |
| 23 | 53.16 | 5.09 | 40.99 | 0.77 |
| 24 | 54.26 | 3.40 | 41.83 | 0.51 |
| 25 | 51.40 | 7.80 | 39.63 | 1.17 |
| 26 | 51.30 | 7.95 | 39.55 | 1.20 |
| 27 | 51.98 | 6.90 | 40.08 | 1.04 |
| 28 | 52.39 | 6.27 | 40.40 | 0.94 |
| 29 | 53.05 | 5.26 | 40.90 | 0.79 |
| 30 | 53.60 | 4.42 | 41.32 | 0.66 |
| 31 | 53.85 | 4.02 | 41.52 | 0.60 |
| 32 | 54.63 | 2.83 | 42.12 | 0.42 |

Table 3: Molar compositions of SM-102 particle library determined by the actual relative flow rates.

| Form. | C12-200 | DOPE | Chol. | PEG |
|---|---|---|---|---|
| 1 | 27.42 | 31.37 | 36.31 | 4.91 |
| 2 | 28.95 | 28.28 | 38.34 | 4.42 |
| 3 | 32.51 | 21.13 | 43.06 | 3.30 |
| 4 | 33.37 | 19.40 | 44.19 | 3.03 |
| 5 | 34.92 | 16.28 | 46.25 | 2.55 |
| 6 | 35.94 | 14.24 | 47.60 | 2.23 |
| 7 | 36.38 | 13.35 | 48.18 | 2.09 |
| 8 | 38.42 | 9.26 | 50.88 | 1.45 |
| 9 | 32.12 | 21.91 | 42.54 | 3.43 |
| 10 | 33.69 | 18.75 | 44.62 | 2.93 |
| 11 | 34.23 | 17.67 | 45.34 | 2.76 |
| 12 | 35.92 | 14.27 | 47.57 | 2.23 |
| 13 | 36.96 | 12.18 | 48.95 | 1.90 |
| 14 | 37.78 | 10.54 | 50.03 | 1.65 |
| 15 | 38.16 | 9.77 | 50.54 | 1.53 |
| 16 | 39.61 | 6.85 | 52.46 | 1.07 |
| 17 | 33.09 | 19.97 | 43.82 | 3.12 |
| 18 | 34.60 | 16.94 | 45.82 | 2.65 |
| 19 | 36.01 | 14.10 | 47.69 | 2.20 |
| 20 | 36.81 | 12.48 | 48.75 | 1.95 |
| 21 | 37.67 | 10.76 | 49.89 | 1.68 |
| 22 | 38.69 | 8.71 | 51.24 | 1.36 |
| 23 | 38.80 | 8.49 | 51.38 | 1.33 |
| 24 | 40.16 | 5.75 | 53.19 | 0.90 |
| 25 | 36.70 | 12.71 | 48.60 | 1.99 |
| 26 | 36.58 | 12.94 | 48.45 | 2.02 |
| 27 | 37.38 | 11.34 | 49.51 | 1.77 |
| 28 | 37.88 | 10.35 | 50.16 | 1.62 |
| 29 | 38.67 | 8.75 | 51.21 | 1.37 |
| 30 | 39.34 | 7.40 | 52.10 | 1.16 |
| 31 | 39.66 | 6.76 | 52.52 | 1.06 |
| 32 | 40.63 | 4.80 | 53.81 | 0.75 |

Table 4: Molar compositions of C12-200 particle library determined by the actual relative flow rates.

Movie S1. Footage of the plate robot moving between wells automatically after initiation. Times in collection positions (in cassettes) and in waste positions were set to 3s each. Pressure regulators on left panel are synced to beginning of robotic motion, rapidly equilibrating between positions. Operating pressures for inputs $i_1$, $i_2$ , $i_3$ (EtOH phase) and $i_4$, $i_5$ , $i_6$ (aqueous phase) are 20, 20, 24, 30, 30, 36 PSI respectively. Regulator communication and timing of movement transition and waste collection were determined from readouts from the Arduino IDE and confirmed using the video footage shown.

### References


[1] Delai Chen, Kevin Love, Yi Chen, Ahmed Eltoukhy, Christian Kastrup, Gaurav Sahay, Alvin Jeon, Yizhou Dong, Kathryn Whitehead, and Daniel Anderson. Rapid discovery of potent sirna-containing lipid nanoparticles enabled by controlled microfluidic formulation. *Journal of the American Chemical Society*, 134:6948–51, 2012. doi: 10.1021/ja301621z. URL `https://www.researchgate.net/profile/Kathryn-Whitehead-3/publication/223956992_Rapid_Discovery_of_Potent_siRNA-Containing_Lipid_Nanoparticles_Enabled_by_Controlled_Microfluidic_Formulation/links/611bcb210c2bfa282a4fb127/Rapid-Discovery-of-Potent-siRNA-Containing-Lipid-Nanoparticles-Enabled-by-Cont pdf`.

[2] Sarah J. Shepherd, Xuexiang Han, Alvin J. Mukalel, Rakan El-Mayta, Ajay S. Thatte, Jingyu Wu, Marshall S. Padilla, Mohamad-Gabriel Alameh, Neha Srikumar, Daeyeon Lee, Drew Weissman, David Issadore, and Michael J. Mitchell. Throughput-scalable manufacturing of sars-cov-2 mrna lipid nanoparticle vaccines. *Proceedings of the National Academy of Sciences*, 120(33):e2303567120, 2023. doi: 10.1073/pnas.2303567120. URL `https://www.pnas.org/doi/full/10.1073/pnas.2303567120`.
